# Genotyping structural variants in pangenome graphs using the vg toolkit

**DOI:** 10.1101/654566

**Authors:** Glenn Hickey, David Heller, Jean Monlong, Jonas A. Sibbesen, Jouni Sirén, Jordan Eizenga, Eric T. Dawson, Erik Garrison, Adam M. Novak, Benedict Paten

**Author notes:** These authors contributed equally to this work (Glenn Hickey, David Heller, Jean Monlong).

## Abstract

Structural variants (SVs) remain challenging to represent and study relative to point mutations despite their demonstrated importance. We show that variation graphs, as implemented in the vg toolkit, provide an effective means for leveraging SV catalogs for short-read SV genotyping experiments. We benchmarked vg against state-of-the-art SV genotypers using three sequence-resolved SV catalogs generated by recent long-read sequencing studies. In addition, we use assemblies from 12 yeast strains to show that graphs constructed directly from aligned *de novo* assemblies improve genotyping compared to graphs built from intermediate SV catalogs in the VCF format.

## Introduction

A structural variant (SV) is a genomic mutation involving 50 or more base pairs. SVs can take several forms such as deletions, insertions, inversions, translocations or other complex events. Due to their greater size, SVs often have a larger impact on phenotype than smaller events such as single nucleotide variants (SNVs) and small insertions and deletions (indels)[1]. Indeed, SVs have long been associated with developmental disorders, cancer and other complex diseases and phenotypes[2].

Despite their importance, SVs remain much more poorly studied than their smaller mutational counterparts. This discrepancy stems from technological limitations. Short read sequencing has provided the basis of most modern genome sequencing studies due to its high base-level accuracy and relatively low cost, but is poorly suited for discovering SVs. The central obstacle is in mapping short reads to the human reference genome. It is generally difficult or impossible to unambiguously map a short read if the sample whose genome is being analyzed differs substantially from the reference at the read’s location. The large size of SVs virtually guarantees that short reads derived from them will not map to the linear reference genome. For example, if a read corresponds to sequence in the middle of a large reference-relative insertion, then there is no location in the reference that corresponds to a correct mapping. The best result a read mapper could hope to produce would be to leave it unmapped. Moreover, SVs often lie in repeat-rich regions, which further frustrate read mapping algorithms.

Short reads can be more effectively used to genotype known SVs. This is important, as even though efforts to catalog SVs with other technologies have been highly successful, their cost currently prohibits their use in large-scale studies that require hundreds or thousands of samples such as disease association studies. Traditional SV genotypers start from reads that were mapped to a reference genome, extracting aberrant mapping that might support the presence of the SV of interest. Current methods such as SVTyper[3] and the genotyping module of Delly[4] (henceforth referred to as Delly Genotyper) typically focus on split reads and paired reads mapped too close or too far from each other. These discordant reads are tallied and remapped to the reference sequence modified with the SV of interest in order to genotype deletions, insertions, duplications, inversions and translocations. SMRT-SV v2 Genotyper uses a different approach: the reference genome is augmented with SV-containing sequences as alternate contigs and the resulting mappings are evaluated with a machine learning model trained for this purpose[5].

The catalog of known SVs in human is quickly expanding. Several large-scale projects have used shortread sequencing and extensive discovery pipelines on large cohorts, compiling catalogs with tens of thousands of SVs in humans[6,7], using split read and discordant pair based methods like Delly[4] to find SVs using short read sequencing. More recent studies using long-read or linked-read sequencing have produced large catalogs of structural variation, the majority of which was novel and sequence-resolved[10,11,5,8,9]. These technologies are also enabling the production of high-quality *de novo* genome assemblies[12,8], and large blocks of haplotype-resolved sequences[13]. Such technical advances promise to expand the amount of known genomic variation in humans in the near future, and further power SV genotyping studies. Representing known structural variation in the wake of increasingly larger datasets poses a considerable challenge, however. VCF, the standard format for representing small variants, is unwieldy when used for SVs due its unsuitability for expressing nested or complex variants. Another strategy consists in incorporating SVs into a linear pangenome reference via alt contigs, but it also has serious drawbacks. Alt contigs tend to increase mapping ambiguity. In addition, it is unclear how to scale this approach as SV catalogs grow.

Pangenomic graph reference representations offer an attractive approach for storing genetic variation of all types[14]. These graphical data structures can seamlessly represent both SVs and point mutations using the same semantics. Moreover, including known variants in the reference makes read mapping, variant calling and genotyping variant-aware. This leads to benefits in terms of accuracy and sensitivity[15,16,17]. The coherency of this model allows different variant types to be called and scored simultaneously in a unified framework.

vg is the first openly available variation graph tool to scale to multi-gigabase genomes. It provides read mapping, variant calling and visualization tools[15]. In addition, vg can build graphs both from variant catalogs in the VCF format and from assembly alignments.

Other tools have used genome graphs or pangenomes to genotype variants. GraphTyper realigns mapped reads to a graph built from known SNVs and short indels using a sliding-window approach[18]. BayesTyper first builds a set of graphs from known variants including SVs, then genotypes variants by comparing the distribution of k-mers in the sequencing reads with the k-mers of haplotype candidate paths in the graph[19]. Paragraph builds a graph for each breakpoint of known variants [20], then, for each breakpoint, it pulls out all nearby reads from the linear alignment and re-aligns them to the graph. Genotypes are computed using the read coverage from the pair of breakpoint graphs corresponding to each SV. These graph-based approaches showed clear advantages over standard methods that use only the linear reference.

In this work, we present a SV genotyping framework based on the variation graph model and implemented in the vg toolkit. We show that this method is capable of genotyping known deletions, insertions and inversions, and that its performance is not inhibited by small errors in the specification of SV allele breakpoints. We evaluated the genotyping accuracy of our approach using simulated and real Illumina reads and a pangenome built from SVs discovered in recent long-read sequencing studies[21,22,23,5], We also compared vg’s performance with state-of-the-art SV genotypers: SVTyper[3], Delly Genotyper[4], BayesTyper[19], Paragraph[20] and SMRT-SV v2 Genotyper[5]. Across the datasets we tested, which range in size from 26k to 97k SVs, vg is the best performing SV genotyper on real short-read data for all SV types in the majority of cases. Finally, we demonstrate that a pangenome graph built from the alignment of *de novo* assemblies of diverse *Saccharomyces cerevisiae* strains improves SV genotyping performance.

## Results

### Structural variation in vg

We used vg to implement a straightforward SV genotyping pipeline. Reads are mapped to the graph and used to compute the read support for each node and edge (see Supplementary Information for a description of the graph formalism). Sites of variation within the graph are then identified using the snarl decomposition as described in [24]. These sites correspond to intervals along the reference paths (ex. contigs or chromosomes) which are embedded in the graph. They also contain nodes and edges deviating from the reference path, which represent variation at the site. For each site, the two most supported paths spanning its interval (haplotypes) are determined, and their relative supports used to produce a genotype at that site (Figure 1a). The pipeline is described in detail in Methods. We rigorously evaluated the accuracy of our method on a variety of datasets, and present these results in the remainder of this section.

**Figure 1:**
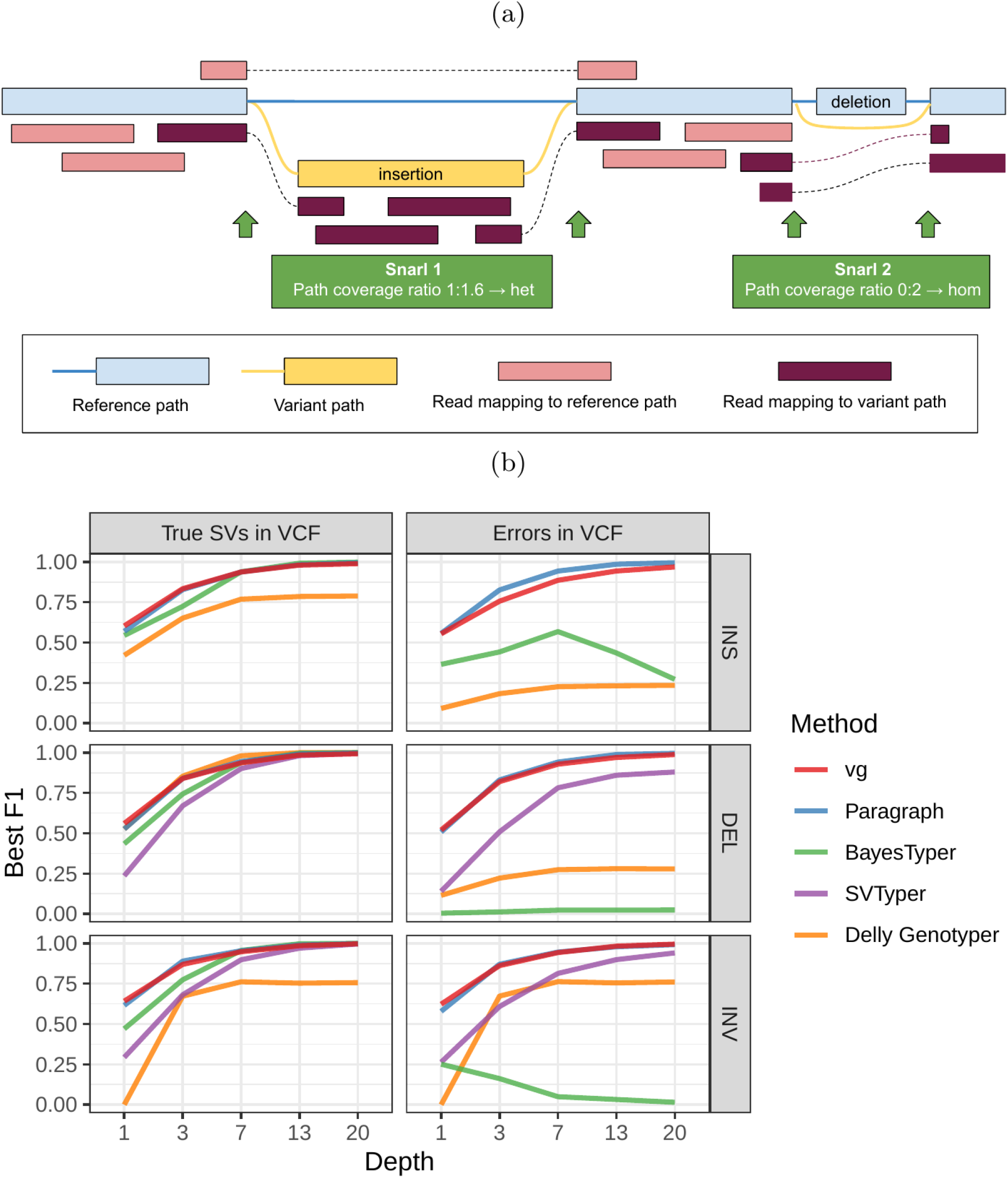
Structural variation in vg. a) vg uses the read coverage over possible paths to genotype variants in a snarl. The cartoon depicts the case of an heterozygous insertion and an homozygous deletion. The algorithm is described in detail in Methods. b) Simulation experiment. Each subplot shows a comparison of genotyping accuracy for five methods. Results are separated between types of variation (insertions, deletions, and inversions). The experiments were also repeated with small random errors introduced to the VCF to simulate breakpoint uncertainty. For each experiment, the x-axis is the simulated read depth and the y-axis shows the maximum F1 across different minimum quality thresholds. SVTyper cannot genotype insertions, hence the missing line in the top panels.

### Simulated dataset

As a proof of concept, we simulated genomes and different types of SVs with a size distribution matching real SVs[22]. We compared vg against Paragraph, SVTyper, Delly Genotyper, and BayesTyper across different levels of sequencing depth. We also added some errors (1-10bp) to the location of the breakpoints to investigate their effect on genotyping accuracy (see Methods). The results are shown in Figure 1b.

When using the correct breakpoints, most methods performed similarly, with differences only becoming visible at very low sequencing depths. Only vg and Paragraph maintained their performance in the presence of 1-10 bp errors in the breakpoint locations. The dramatic drop for BayesTyper can be explained by its k-mer-based approach that requires precise breakpoints. Overall, these results show that vg is capable of genotyping SVs and is robust to breakpoint inaccuracies in the input VCF.

### HGSVC dataset

72,485 structural variants from The Human Genome Structural Variation Consortium (HGSVC) were used to benchmark the genotyping performance of vg against the four other SV genotyping methods. This high-quality SV catalog was generated from three samples using a consensus from different sequencing, phasing, and variant calling technologies[22]. The three individual samples represent different human populations: Han Chinese (HG00514), Puerto-Rican (HG00733), and Yoruban Nigerian (NA19240). We used these SVs to construct a graph with vg and as input for the other genotypers. Using short sequencing reads, the SVs were genotyped and compared with the genotypes in the original catalog (see Methods).

First we compared the methods using simulated reads for HG00514. This represents the ideal situation where the SV catalog exactly matches the SVs supported by the reads. BayesTyper and vg showed the best F1 score and precision-recall trade-offs (Figures 2a and S1, Table S1), outperforming the other methods by a clear margin. When restricting the comparisons to regions not identified as tandem repeats or segmental duplications, the genotyping predictions were significantly better for all methods. We observed similar results when evaluating the presence of an SV call instead of the exact genotype (Figures 2a and S2).

**Figure 2:**
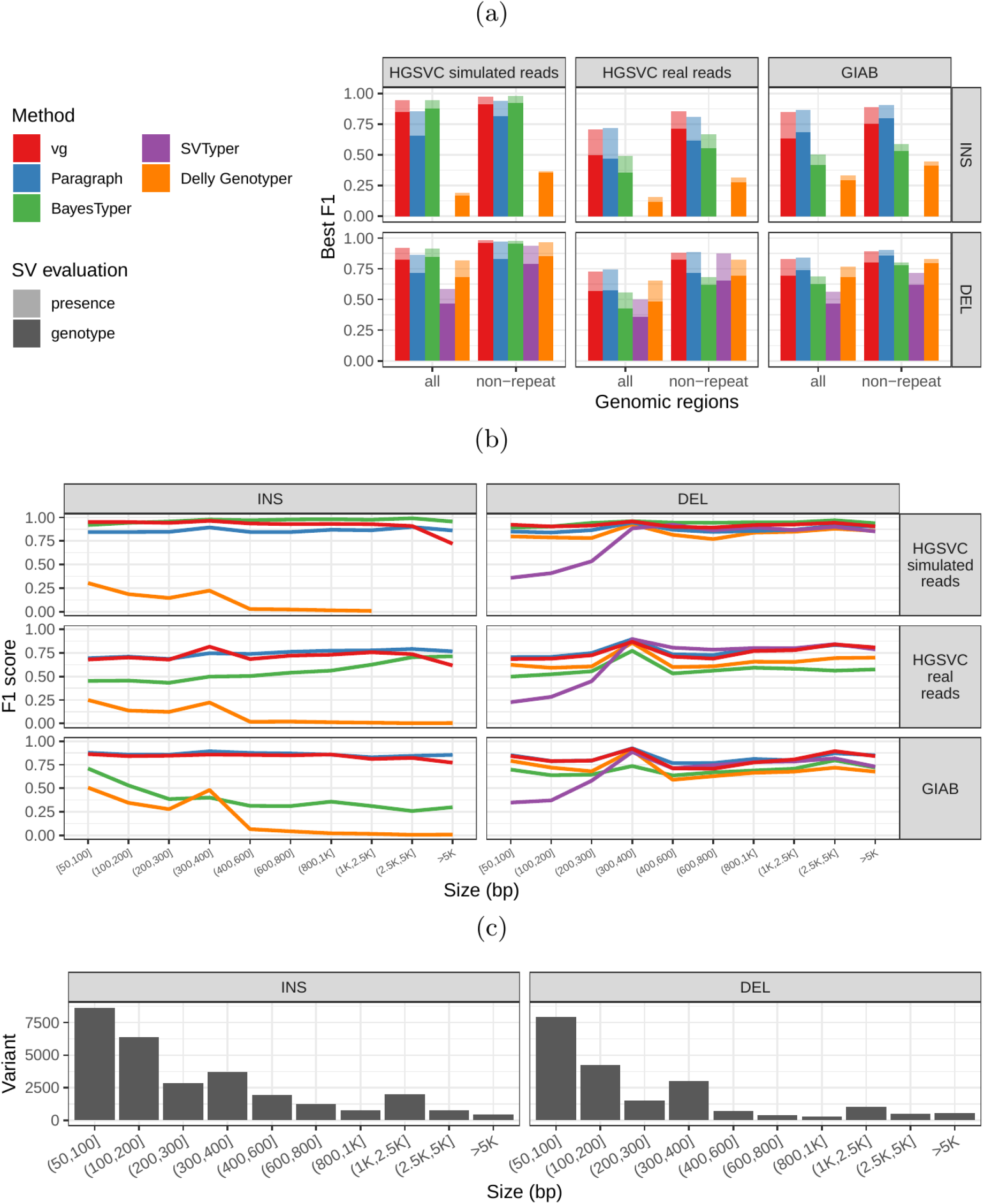
Structural variants from the HGSVC and Genome in a Bottle datasets. HGSVC: Simulated and real reads were used to genotype SVs and compared with the high-quality calls from Chaisson et al.[22]. Reads were simulated from the HG00514 individual. Using real reads, the three HG00514, HG00733, and NA19240 individuals were tested. GIAB: Real reads from the HG002 individual were used to genotype SVs and compared with the high-quality calls from the Genome in a Bottle consortium[21,23,25]. a) Maximum F1 score for each method (color), across the whole genome or focusing on non-repeat regions (x-axis). We evaluated the ability to predict the presence of an SV (transparent bars) and the exact genotype (solid bars). Results are separated across panels by variant type: insertions and deletions. SVTyper cannot genotype insertions, hence the missing bars in the top panels. b) Maximum F1 score for different size classes when evaluating on the presence of SVs across the whole genome. c) Size distribution of SVs in the HGSVC and GIAB catalogs.

We then repeated the analysis using real Illumina reads from the three HGSVC samples to benchmark the methods on a more realistic experiment. Here, vg clearly outperformed other approaches (Figures 2a and S3). In non-repeat regions and insertions across the whole genome, the F1 scores and precision-recall AUC were higher for vg compared to other methods. For example, for deletions in non-repeat regions, the F1 score for vg was 0.824 while the second best method, Paragraph, had a F1 score of 0.717. We observed similar results when evaluating the presence of an SV call instead of the exact genotype (Figures 2a and S4).

In general, the genotyped variants were matched 1-to-1 with variants in the truth set but some methods showed some signs of “over-genotyping” that is not reflected in the precision/recall/F1 scores. Methods like Paragraph, Delly Genotyper or SVTyper tended to genotype on average more than one variant per truth-set variant (Figure S5). Like other SV catalogs, the HGSVC catalog is not fully sequence-resolved and contains a number of near-duplicates with slightly different breakpoint deflnition. When genotyping a sample, multiple versions of a variant are genotyped multiple times by methods that analyze each variant independently. In contrast, vg follows a unified path-centric approach that only select the best genotype in a region (see Methods).

We further evaluate the performance for different SV sizes and repeat content. In addition, vg’s performance was stable across the spectrum of SV sizes (Figure 2b-c). By annotating the repeat content of the deleted/inserted sequence we further evaluated vg’s performance across repeat classes. As expected, simple repeat variation was more challenging to genotype than transposable element polymorphisms (Figure S6). Figure 3 shows an example of an exonic deletion that was correctly genotyped by vg but not by BayesTyper, SVTyper or Delly Genotyper.

**Figure 3:**
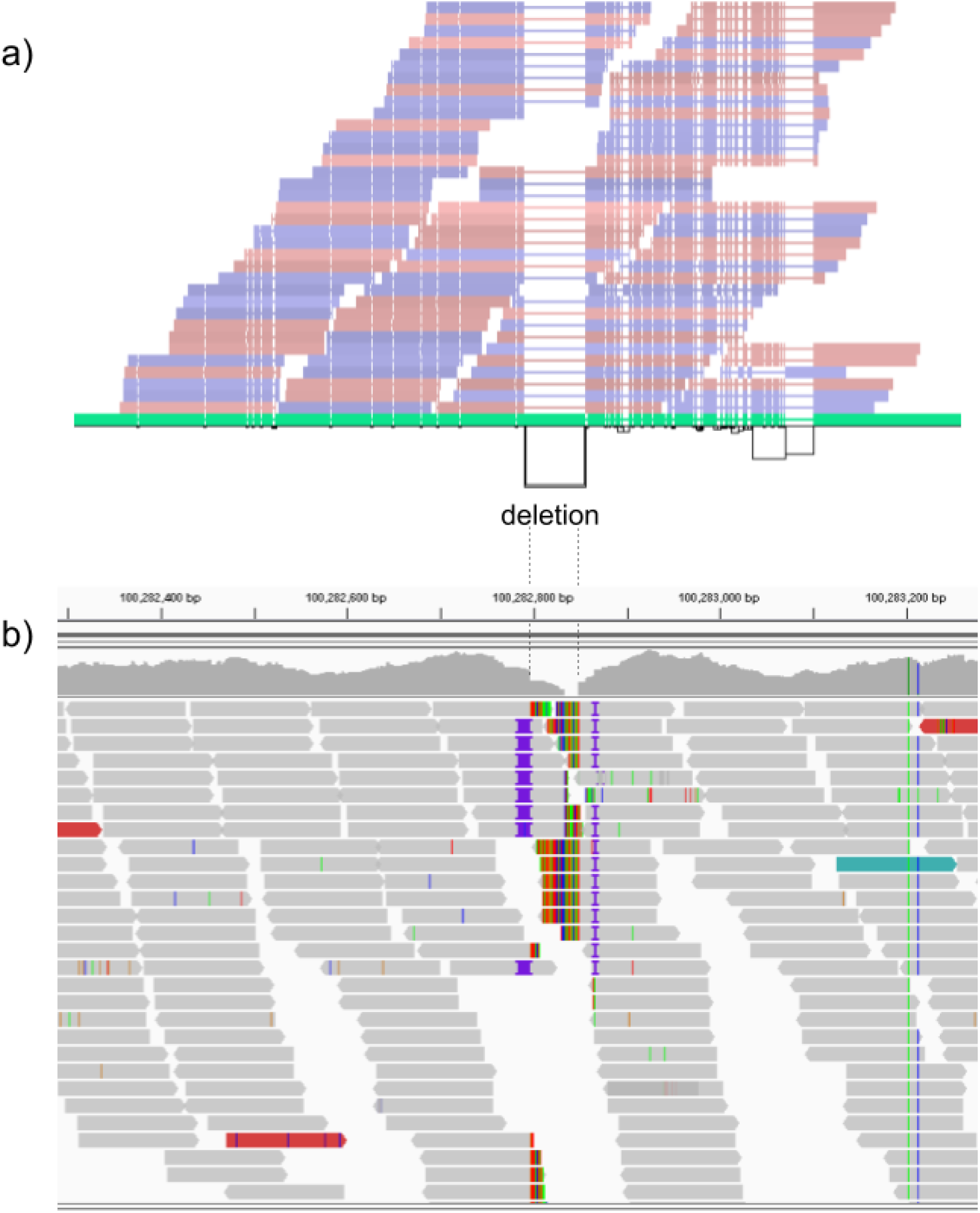
Exonic deletion in the HGSVC dataset correctly genotyped by vg. a) Visualization of the HGSVC graph as augmented by reads aligned by vg at a locus harboring a 51 bp homozygous deletion in the UTR region of the LONRF2 gene. At the bottom, a horizontal black line represents the topologically sorted nodes of the graph. Black rectangles represent edges found in the graph. Above this rendering of the topology, the reference path from GRCh38 is shown (in green). Red and blue bars represent reads mapped to the graph. Thin lines in the reference path and read mappings highlight relative gaps (either insertions or deletions) against the full graph. The vg read mappings show consistent coverage even over the deletion. b) Reads mapped to the linear genome reference GRCh38 using bwa mem[26] in the same region. Reads contain soft-clipped sequences and short insertions near the deletion breakpoints. Part of the deleted region is also covered by several reads, potentially confusing traditional SV genotypers.

### Other long-read datasets

#### Genome in a Bottle Consortium

The Genome in a Bottle (GiaB) consortium is currently producing a high-quality SV catalog for an Ashkenazim individual (HG002)[21,23,25]. Dozens of SV callers operating on datasets from short, long, and linked reads were used to produce this set of SVs. We evaluated the SV genotyping methods on this sample as well using the GIAB VCF, which also contains parental calls (HG003 and HG004), all totaling 30,224 SVs. Relative to the HGSVC dataset, vg performed similarly but Paragraph saw a large boost in accuracy and was the most accurate method across all metrics. (Figures 2, S7 and S8, and Table S2). As before, the remaining methods produced lower F1 scores.

#### SMRT-SV v2 catalog and training data [5]

A recent study by Audano et al. generated a catalog of 97,368 SVs (referred as SVPOP below) using long-read sequencing across 15 individuals[5]. These variants were then genotyped from short reads across 440 individuals using the SMRT-SV v2 Genotyper, a machine learning-based tool implemented for that study. The SMRT-SV v2 Genotyper was trained on a pseudo-diploid genome constructed from high quality assemblies of two haploid cell lines (CHM1 and CHM13) and a single negative control (NA19240). We first used vg to genotype the SVs in this two-sample training dataset using 30X coverage reads, and compared the results with the SMRT-SV v2 Genotyper. vg was systematically better at predicting the presence of an SV for both SV types, but SMRT-SV v2 Genotyper produced slightly better genotypes for deletions in the whole genome(see Figures 4, S9 and S10, and Table S3). To compare vg and SMRT-SV v2 Genotyper on a larger dataset, we then genotyped SVs from the entire SVPOP catalog with both methods, using the read data from the three HGSVC samples described above. Given that the SVPOP catalog contains these three samples, we once again evaluated accuracy by using the long-read calls as a baseline. Paragraph was included as an additional point of comparison.

**Figure 4:**
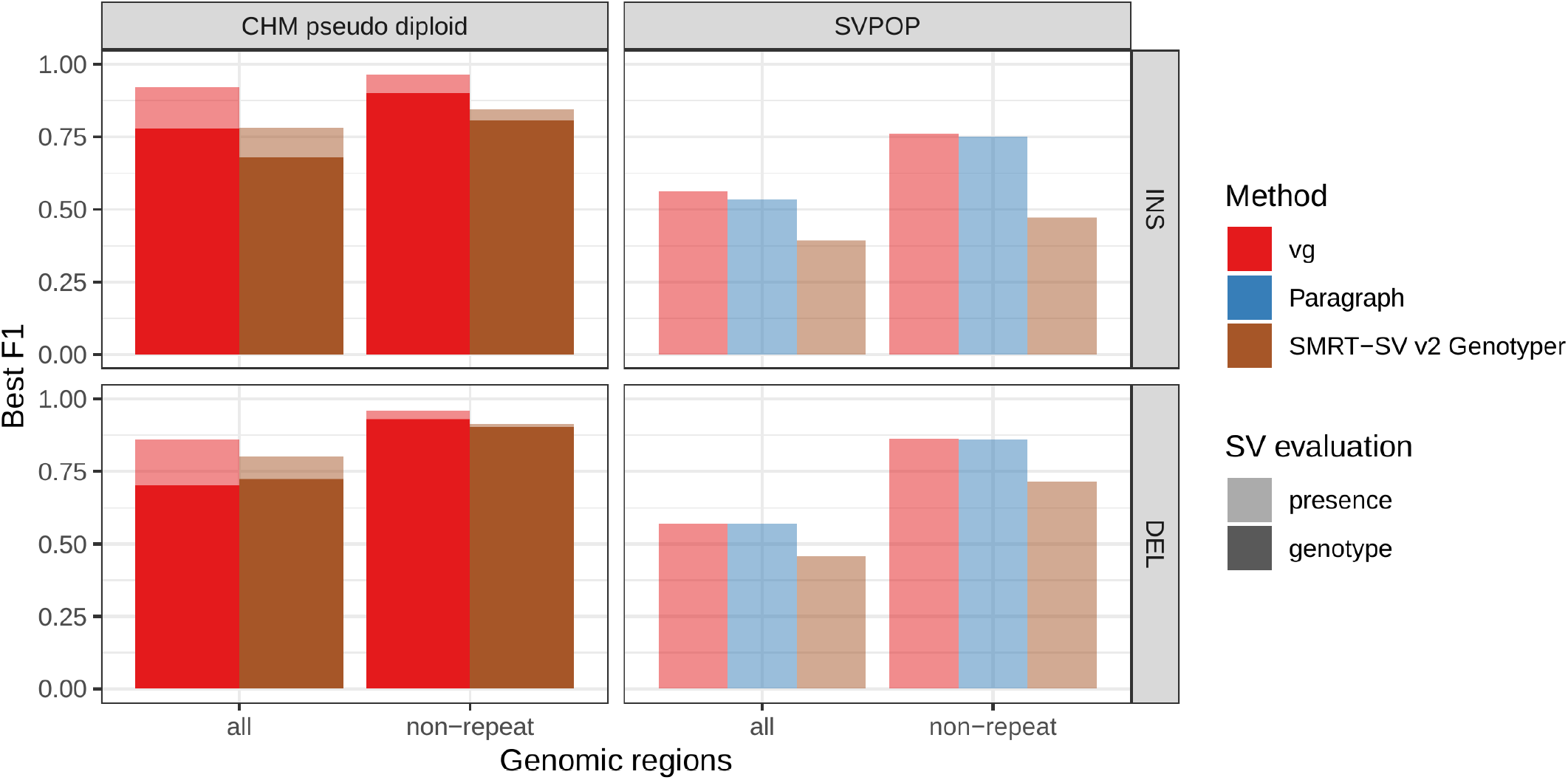
Structural variants from SMRT-SV v2 [5]. The pseudo-diploid genome built from two CHM cell lines and one negative control sample was originally used to train SMRT-SV v2 Genotyper in Audano et al.[5]. It contains 16,180 SVs. The SVPOP panel shows the combined results for the HG00514, HG00733, and NA19240 individuals, three of the 15 individuals used to generate the high-quality SV catalog in Audano et al. [5]. Here, we report the maximum F1 score (y-axis) for each method (color), across the whole genome or focusing on non-repeat regions (x-axis). We evaluated the ability to predict the presence of an SV (transparent bars) and the exact genotype (solid bars). Genotype information is not available in the SVPOP catalog hence genotyping performance could not be evaluated.

Compared to SMRT-SV v2 Genotyper, vg had a better precision-recall curve and a higher F1 for both insertions and deletions (SVPOP in Figures 4 and S11, and Table S4). Paragraph’s performance was virtually identical to vg’s. Of note, SMRT-SV v2 Genotyper produces *no-calls* in regions where the read coverage is too low, and we observed that its recall increased when filtering these regions out the input set. Interestingly, vg performed well even in regions where SMRT-SV v2 Genotyper produced *nocalls* (Figure S12 and Table S5). Audano et al. discovered 217 sequence-resolved inversions using long reads, which we attempted to genotype. vg correctly predicted the presence of around 14% of the inversions present in the three samples (Table S4). Inversions are often complex, harboring additional variation that makes their characterization and genotyping challenging.

### Graphs from alignment of *de novo* assemblies

We can construct variation graphs directly from whole genome alignments (WGA) of multiple *de novo* assemblies[15]. This bypasses the need for generating an explicit variant catalog relative to a linear reference, which could be a source of error due to the reference bias inherent in read mapping and variant calling. Genome alignments from graph-based software such as Cactus [27] can contain complex structural variation that is extremely difficult to represent, let alone call, outside of a graph, but which is nevertheless representative of the actual genomic variation between the aligned assemblies. We sought to establish if graphs built in this fashion provide advantages for SV genotyping.

To do so, we analyzed public sequencing datasets for 12 yeast strains from two related clades (*S. cerevisiae* and *S. paradoxus)* [28]. We distinguished two different strain sets, in order to assess how the completeness of the graph affects the results. For the *all strains set*, all 12 strains were used, with *S.c. S288C* as the reference strain. For the *five strains set, S.c. S288C* was used as the reference strain, and we selected two other strains from each of the two clades (see Methods). We compared genotyping results from a WGA-derived graph (*cactus graph*) with results from a VCF-derived graph (*VCF graph*). The *VCF graph* was created from the linear reference genome of the *S.c. S288C* strain and a set of SVs relative to this reference strain in VCF format identified from the other assemblies in the respective strain set by three methods: Assemblytics [29], AsmVar [30] and paftools [31]. The *cactus graph* was derived from a multiple genome alignment of the strains in the respective strain set using Cactus [27]. The *VCF graph* is mostly linear and highly dependent on the reference genome. In contrast, the *cactus graph* is structurally complex and relatively free of reference bias.

First, we tested our hypothesis that the *cactus graph* has higher mappability due to its better representation of sequence diversity among the yeast strains (see Supplementary Information). Generally, more reads mapped to the *cactus graph* with high identity (Figures S13a and S14a) and high mapping quality (Figures S13b and S14b) than to the *VCF graph*. On average, 88%, 79%, and 68% of reads mapped to the *all strain cactus graph* with an identity of at least 50%, 90%, and 100%, respectively, compared to only 77%, 57%, and 23% of reads on the *all strain VCF graph*. Similarly, 88% of reads mapped to the *all strain cactus graph* with a mapping quality of at least 30 compared to only 80% of reads on the *all strain VCF graph*.

Next, we compared the SV genotyping performance of both graph types. We mapped short reads from the 11 non-reference strains to both graphs and genotyped variants for each strain using the vg toolkit’s variant calling module (see Methods). There is no gold standard available for these samples to compare against which renders an evaluation using recall, precision and F1 score impossible. Therefore, we used an indirect measure of SV genotyping accuracy. We evaluated each SV genotype set based on the alignment of reads to a *sample graph* constructed from the genotype set (see Methods). Conceptually, the sample graph represents the sample’s diploid genome by starting out from the reference genome and augmenting it with the genotype results. If a given genotype set is correct, we expect that reads from the same sample will be mapped with high identity and confidence to the corresponding sample graph. To specifically quantify mappability in SV regions we excluded reads that produced identical mapping quality and identity on both sample graphs and an empty sample graph containing the linear reference only (see Methods and Figure S15 for results from all reads). Then, we analyzed the average delta in mapping identity and mapping quality of the remaining short reads between both sample graphs (Figures 5a and b).

**Figure 5:**
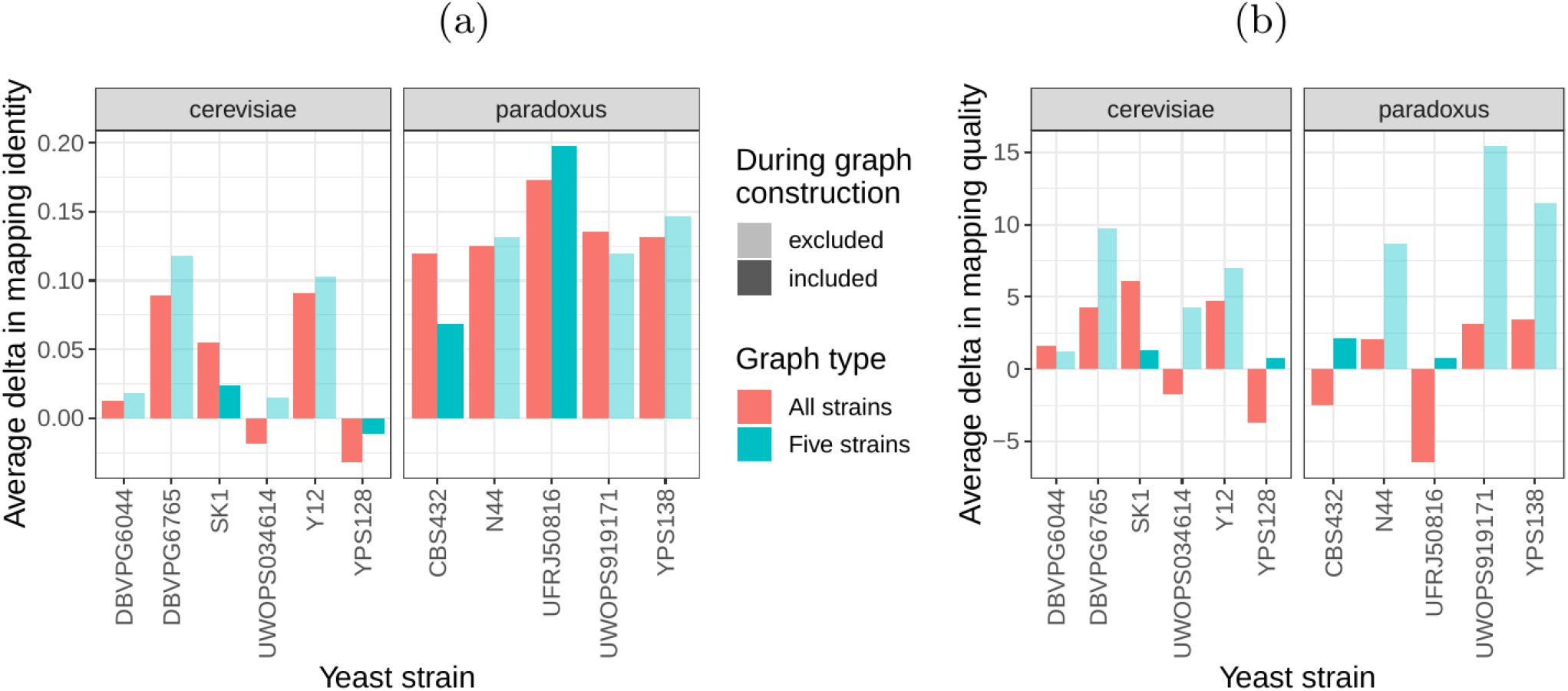
SV genotyping comparison. Short reads from all 11 non-reference yeast strains were used to genotype SVs contained in the *cactus graph* and the *VCF graph*. Subsequently, sample graphs were generated from the resulting SV genotype sets. The short reads were aligned to the sample graphs and reads with identical mapping identity and quality across both sample graphs and an additional empty sample graph were removed from the analysis. The quality of the remaining divergent alignments was used to ascertain SV genotyping performance. The bars show the average delta in mapping identity (a) and in mapping quality (b) of divergent short reads aligned to the sample graphs derived from the *cactus graph* and the *VCF graph*. Positive values denote an improvement of the *cactus graph* over the *VCF graph*. Colors represent the two strain sets and transparency indicates whether the respective strain was part of the *five strains set*.

For most of the strains, we observed an improvement in mapping identity of the short reads on the *cactus sample graph* compared to the *VCF sample graph*. The mean improvement in mapping identity across the strains (for reads differing in mapping identity) was 8.0% and 8.5% for the *all strains set* graphs and the *five strains set* graphs, respectively. Generally, the improvement in mapping identity was larger for strains in the *S. paradoxus* clade (mean of 13.7% and 13.3% for the two strain sets, respectively) than for strains in the *S. cerevisiae* clade (mean of 3.3% and 4.4%). While the higher mapping identity indicated that the *cactus graph* represents the reads better (Figure 5a), the higher mapping quality confirmed that this did not come at the cost of added ambiguity or a more complex graph (Figure 5b). For most strains, we observed an improvement in mapping quality of the short reads on the *cactus sample graph* compared to the *VCF sample graph* (mean improvement across the strains of 1.0 and 5.7 for the two strain sets, respectively).

## Discussion

Overall, graph-based methods were more accurate than traditional SV genotypers in our benchmarks, with vg performing best across most datasets. These results show that SV genotyping benefits from variant-aware read mapping and graph based genotyping, a finding consistent with previous studies[15,16,17,18,19]. Paragraph, another graph-based genotyper which was released as we were submitting this work, was very competitive with vg and showed the best overall accuracy on the GIAB dataset. In addition to being featured prominently in Paragraph’s development and evaluation, the GIAB dataset we used was a different coverage (50X) than the other 30X datasets we used. Our simulation results show that Paragraph is slightly more robust than vg with respect to differences in coverage and perhaps this is a factor in the difference in performance. In the future, we would like to better model the expected read depth in the vg genotyper as it currently does not exploit this information. In contrast, vg is much more accurate than Paragraph on the HGSVC dataset and we speculate that this is due to the higher number of overlapping variants. Using the snarl decomposition, vg can genotype arbitrary combinations of SVs simultaneously, whereas Paragraph operates one at a time.

We took advantage of newly released datasets for our evaluation, which feature up to 3.7 times more variants than the more widely-used GIAB benchmark. More and more large-scale projects are using low cost short-read technologies to sequence the genomes of thousands to hundreds of thousands of individuals (e.g. the Pancancer Analysis of Whole Genomes[32], the Genomics England initiative[33], and the TOPMed consortium[34]). We believe pangenome graph-based approaches will improve both how efficiently SVs can be represented, and how accurately they can be genotyped with this type of data.

A particular advantage of our method is that it does not require exact breakpoint resolution in the variant library. Our simulations showed that vg’s SV genotyping algorithm is robust to errors of as much as 10 bp in breakpoint location. However, there is an upper limit to this flexibility, and we find that vg cannot accurately genotype variants with much higher uncertainty in the breakpoint location (like those discovered through read coverage analysis). vg is also capable of fine-tuning SV breakpoints by augmenting the graph with differences observed in read alignments. Simulations showed that this approach can usually correct small errors in SV breakpoints (Figure S16 and Table S6).

vg uses a unified framework to call and score different variant types simultaneously. In this work, we only considered graphs containing certain types of SVs, but the same methods can be extended to a broader range of graphs. For example, we are interested in evaluating how genotyping SVs together with SNPs and small indels using a combined graph effects the accuracy of studying either alone. The same methods used for genotyping known variants in this work can also be extended to call novel variants by first augmenting the graph with edits from the mapped reads. This approach, which was used only in the breakpoint fine-tuning portion of this work, could be further used to study small variants around and nested within SVs. Novel SVs could be called by augmenting the graph with long-read mappings. vg is entirely open source, and its ongoing development is supported by a growing community of researchers and users with common interest in scalable, unbiased pangenomic analyses and representation. We expect this collaboration to continue to foster increases in the speed, accuracy and applicability of methods based on pangenome graphs in the years ahead.

Our results suggest that constructing a graph from *de novo* assembly alignment instead of a VCF leads to better SV genotyping. High quality *de novo* assemblies for human are becoming more and more common due to improvements in technologies like optimized mate-pair libraries[35] and long-read sequencing[12]. We expect future graphs to be built from the alignment of numerous *de novo* assemblies, and we are presently working on scaling our assembly-based pipeline to human-sized genome assemblies. Another challenge is creating genome graphs that integrate assemblies with variant-based data resources. One possible approach is to progressively align assembled contigs into variation graphs constructed from variant libraries, but methods for doing so are still experimental.

## Conclusion

In this study, the vg toolkit was compared to existing SV genotypers across several high-quality SV catalogs. We showed that its method of mapping reads to a variation graph leads to better SV genotyping compared to other state-of-the-art methods. This work introduces a flexible strategy to integrate the growing number of SVs being discovered with higher resolution technologies into a unifled framework for genome inference. Our work on whole genome alignment graphs shows the benefit of directly utilizing *de novo* assemblies rather than variant catalogs to integrate SVs in genome graphs. We expect this latter approach to increase in significance as the reduction in long read sequencing costs drives the creation of numerous new *de novo* assemblies. We envision a future in which the lines between variant calling, genotyping, alignment, and assembly are blurred by rapid changes in sequencing technology. Fully graph based approaches, like the one we present here, will be of great utility in this new phase of genome inference.

## Methods

### SV Genotyping Algorithm

The input to the SV genotyping algorithm is an indexed variation graph in `xg` format along with a (single-sample) read alignment in `GAM` format. If the graph was constructed from a VCF, as was the case for the human-genome graphs discussed in this paper, this VCF can also be input to the caller. The first step is to compute a compressed coverage index from the alignment using this command, `vg pack <graph.xg> <alignment.gam> –Q 5 –o graph.pack`. This index stores the number of reads with mapping quality at least 5 mapped to each edge and each base of each node on the graph. Computing the coverage can be done in a single scan through the reads and, in practice, tends to be an order of magnitude faster than sorting the reads.

Variation graphs, as represented in vg, are bidirected. In a bidirected graph, every node can be thought of having two distinct *sides*. See, for example, the left and right sides of each rectangle in Figure 1a. If *x* is the side of a given node *A*, then we use the notation *x’* to denote the other side of *A*. A snarl is defined by a pair of sides, *x* and *y*, that satisfy the following criteria:

1. Removing all edges incident to *x’* and *y’* disconnects the graph, creating a connected component *X* that contains *x* and *y*.
2. There is no side *z* in *X* such that *{x,z}* satisfies the above criteria. Likewise for *y*.

Snarls can be computed in linear time using a cactus graph decomposition [24]. They can be computed once for a given graph using `vg snarls`, or on the fly with `vg call`.

Once the snarls have been identified, the SV genotyping algorithm proceeds as follows. For every snarl in the graph for which both end nodes lie on a reference path (such as a chromosome) and that it is not contained in another snarl, the following steps are performed.

1. All VCF variants, *v1, v2,…, vk* that are contained within the snarl are looked up using information embedded during graph construction. Let *|vi|* be the number of alleles in the *ith* VCF variant. Then there are *|v1|*x*|v2|*…x*|vk|* possible haplotypes through the snarl. If this number is too high (>500,000), then alleles with average support of less than 1 are filtered out.
2. For each possible haplotype, a corresponding bidrected path through the snarl (from *x* to *y*) is computed.
3. For each haplotype path, its average support (over bases and edges) is computed using the compressed coverage index, and the two most-supported paths are selected (ties are broken arbitrarily).
4. If the most supported path exceeds the minimum support threshold (default 1), and has more than *B* (default 6) times the support of the next most supported path, the site is called homozygous for the allele associated with the most supported path.
5. Else if the second most supported path exceeds the minimum support threshold (default 1), then the site is deemed heterozygous with an allele from each of the top two paths.
6. Given the genotype computed above, it is trivial to map back from the chosen paths to the VCF alleles in order to produce the final output.

The command to do the above is vg call `<graph.xg> –k <graph.pack> –v variants.vcf.gz` If the graph was not constructed from a VCF, then a similar algorithm is used except the traversals are computed heuristically searching through the graph. This is enabled by not using the `–v` option in the above command.

#### toil-vg

toil-vg is a set of Python scripts for simplifying vg tasks such as graph construction, read mapping and SV genotyping. Much of the analysis in this report was done using toil-vg, with the exact commands available at github.com/vgteam/sv-genotyping-paper. toil-vg uses the Toil workflow engine [36] to seamlessly run pipelines locally, on clusters or on the cloud. Graph indexing, and mapping in particular are computationally expensive (though work is underway to address this) and well-suited to distribution on the cloud. The principal toil-vg commands used are described below.

#### toil-vg construct

toil-vg construct automates graph construction and indexing following the best practices put forth by the vg community. Graph construction is parallelized across different sequences from the reference FASTA, and different whole-genome indexes are created side by side when possible. The graph is automatically annotated with paths corresponding to the different alleles in the input VCF. The indexes created are the following:

- xg index: This is a compressed version of the graph that allows fast node, edge and path lookups
- gcsa2 index: This is a substring index used only for read mapping
- gbwt index: This is an index of all the haplotypes in the VCF as implied by phasing information. When available, it is used to help ensure that haplotype information is preserved when constructing the gcsa2 index
- snarls index: The snarls represent sites of variation in the graph and are used for genotyping and variant calling.

#### toil-vg map

toil-vg map splits the input reads into batches, maps each batch in parallel, then merges the result.

#### toil-vg call

toil-vg call splits the input graph by chromosome and calls each one individually. `vg call` has been recently updated so that this subdivision is largely unnecessary: the entire graph can be easily called at once. Still, toil-vg can be used to farm this task out to a single cloud node if desired.

#### toil-vg sveval

toil-vg sveval evaluates the SV calls relative to a truth set. Matching SV calls is non-trivial because two SV callsets often differs slightly around the breakpoints. Even for a genotyping experiment, the same input SVs can have equivalent but different representations. Furthermore, SV catalogs often contain very similar SVs that could be potentially duplicates of the same true variant. To make sure that SVs are matched properly when comparing genotyped SVs and the truth set, we use an approach that overlaps variants and aligns allelic sequences if necessary. It was implemented in the sveval R package (https://github.com/jmonlong/sveval). Figure S17 shows an overview of the SV evaluation approach which is described below. Of note, the variants are first normalized with `bcftools norm` (1.9) to ensure consistent representation between called variants and baseline variants[37].

For deletions and inversions, we begin by computing the overlaps between the SVs in the call set and the truth set. For each variant we then compute the proportion of its region that is covered by a variant in the other set, considering only variants overlapping with at least 10% reciprocal overlap. If this coverage proportion is higher than 50%, we consider the variant *covered*. True positives (TPs) are covered variants from the call set (when computing the precision) or the truth set (when computing the recall). Variants from the call set are considered false positives (FPs) if they are not covered by the truth set. Conversely, variants from the truth set are considered false negatives (FNs) if they are not covered by the call set.

For insertions, we select pairs of insertions that are located no farther than 20 bp from each other. We then align the inserted sequences using a Smith-Waterman alignment. For each insertion we compute the proportion of its inserted sequence that aligns a matched variant in the other set. If this proportion is at least 50% the insertions are considered covered. Covering relationships are used to define TPs, FPs, and FNs the same way as for deletions and inversions.

The results shown in this study used a minimum of 50% coverage to match variants but we also replicated the results using 90% minimum coverage and observed similar results (see Figure S18).

The coverage statistics are computed using any variant larger than 1 bp but a minimum size is required for a variant to be counted as TP, FP, or FN. In this work, we used the default minimum SV size of 50 bp.

sveval accepts VCF files with symbolic or explicit representation of the SVs. If the explicit representation is used, multi-allelic variants are split and their sequences right-trimmed. When using the explicit representation and when the REF and ALT sequences are longer than 10 bp, the reversecomplement of the ALT sequence is aligned to the REF sequence to identify potential inversions. If more than 80% of the sequence aligns, it is classified as an inversion.

We assess both the ability to predict the presence of an SV and the full genotype. For the *presence* evaluation, both heterozygous and homozygous alternate SVs are compared jointly using the approach described above. To compute genotype-level metrics, the heterozygous and homozygous SVs are compared separately. Before splitting the variants by genotype, pairs of heterozygous variants with reciprocal overlap of at least 80% are merged into a homozygous ALT variant. To handle fragmented variants, consecutive heterozygous variants located at less that 20 bp from each other are first merged into larger heterozygous variants.

Precision-recall curves are produced by successively filtering out variants of low-quality. By default, the *QUAL* field in the VCF file is used as the quality information. If *QUAL* is missing (or contains only 0s), the genotype quality in the *GQ* field is used.

The evaluation is performed using all variants or using only variants within high-confidence regions. In most analysis, the high-confidence regions are constructed by excluding segmental duplications and tandem repeats (using the respective tracks from the UCSC Genome Browser). For the GIAB analysis, we used the Tier 1 high-confidence regions provided by the GIAB consortium in version 0.6.

The inserted/deleted sequence was also annotated using RepeatMasker[38]. SVs were separated by repeat family if the annotated repeat element covered more than 80% of the sequence. We recomputed precision and recall in the most frequent repeat families.

The average number of genotyped variants per variant in the truth set (Figure S5) was computed by dividing the number of TPs from the call set by the number of TPs from the truth set, i.e. the ratio of matched variants between the two variant sets.

### Other SV genotypers

#### BayesTyper (v1.5 beta 62888d6)

Where not specified otherwise BayesTyper was run as follows. Raw reads were mapped to the reference genome using `bwa mem` [26] (0.7.17). GATK haplotypecaller[39] (3.8) and Platypus[40] (0.8.1.1) with assembly enabled were run on the mapped reads to call SNVs and short indels (<50bp) needed by BayesTyper for correct genotyping. The VCFs with these variants were then normalized using `bcftools norm` (1.9) and combined with the SVs across samples using `bayesTyperTools combine` to produce the input candidate set. k-mers in the raw reads were counted using kmc[41] (3.1.1) with a k-mer size of 55. A Bloom filter was constructed from these k-mers using `bayesTyperTools makeBloom`. Finally, variants were clustered and genotyped using `bayestyper cluster` and `bayestyper genotype`, respectively, with default parameters except `--min–genotype–posterior 0`. Non-PASS variants and non-SVs (GATK and Platypus origin) were filtered prior to evaluation using `bcftools filter` and `filterAlleleCallsetOrigin`, respectively.

#### Delly (v0.7.9)

The `delly call` command was run on the reads mapped by `bwa mem` [26], the reference genome FASTA file, and the VCF containing the SVs to genotype (converted to their explicit representations).

#### SVTyper (v0.7.0)

The VCF containing deletions was converted to symbolic representation and passed to `svtyper` with the reads mapped by `bwa mem` [26]. The output VCF was converted back to explicit representation using `bayesTyperTools convertAllele` to facilitate variant normalization before evaluation.

#### Paragraph (v2.3)

Paragraph was run using default parameters using the `multigrmpy.py` script, taking the input VCF and reads mapped by `bwa mem` [26] as inputs. We used the genotype estimates in the `genotypes.vcf.gz` output file. In order for Paragraph to run, we added padding sequence to problematic variants in the input VCFs of the GIAB and SVPOP catalogs.

#### SMRT-SV v2 Genotyper (v2.0.0 Feb 21 2019 commit adb13f2)

SMRT-SV v2 Genotyper was run with the “30x-4” model and min-call-depth 8 cutoff. It was run only on VCFs created by SMRT-SV, for which the required contig BAMs were available. The Illumina BAMs used where the same as the other methods described above. The output VCF was converted back to explicit representation to facilitate variant normalization later.

#### Running time and memory usage

Running times and memory usage for the different tools are shown in Table S7. The *Elapsed (wall clock) time* and the *Maximum resident set size* were extracted from the output of `/usr/bin/time – v`. We show the profiling results when genotyping the HGSVC SV catalog in the HG00514 sample.

### Simulation experiment

We simulated a synthetic genome with 1000 insertions, deletions and inversions. We separated each variant from the next by a buffer of at least 500 bp. The sizes of deletions and insertions followed the distribution of SV sizes from the HGSVC catalog. We used the same size distribution as deletions for inversions. A VCF file was produced for three simulated samples with genotypes chosen uniformly between homozygous reference, heterozygous, and homozygous alternate.

We created another VCF file containing errors in the SV breakpoint locations. We shifted one or both breakpoints of deletions and inversions by distances between 1 and 10 bp. The locations and sequences of insertions were also modified, either shifting the variants or shortening them at the flanks, again by up to 10 bp.

Paired-end reads were simulated using `vg sim` on the graph that contained the true SVs. Different read depths were tested: 1x, 3x, 7x, 10x, 13x, 20x. The base qualities and sequencing errors were trained to resemble real Illumina reads from NA12878 provided by the Genome in a Bottle Consortium.

The genotypes called in each experiment (genotyping method/VCF with or without errors/sequencing depth) were compared to the true SV genotypes to compute the precision, recall and F1 score (see toil-vg sveval).

#### Breakpoint fine-tuning using graph augmentation

vg can call variants after augmenting the graph with the read alignments to discover new variants (see toil-vg call). We tested if this approach could fine-tune the breakpoint location of SVs in the graph. We started with the graph that contained approximate SVs (1-10 bp errors in breakpoint location) and 20x simulated reads from the simulation experiment (see Simulation experiment). The variants called after graph augmentation were compared with the true SVs. We considered fine-tuning correct if the breakpoints matched exactly.

### HGSVC Analysis

We first obtained phased VCFs for the three Human Genome Structural Variation Consortium (HGSVC) samples from Chaisson et al.[22] and combined them with `bcftools merge`. A variation graph was created and indexed using the combined VCF and the HS38D1 reference with alt loci excluded. The phasing information was used to construct a GBWT index[42], from which the two haploid sequences from HG00514 were extracted as a graph. Illumina read pairs with 30x coverage were simulated from these sequences using vg sim, with an error model learned from real reads from the same sample. These simulated reads reflect an idealized situation where the breakpoints of the SVs being genotyped are exactly known *a priori*. The reads were mapped to the graph, and the mappings used to genotype the SVs in the graph. Finally, the SV calls were compared back to the HG00514 genotypes from the HGSVC VCF. We repeated the process with the same reads on the linear reference, using `bwa mem` [26] for mapping and Delly Genotyper, SVTyper, Paragraph and BayesTyper for SV genotyping.

We downloaded Illumina HiSeq 2500 paired end reads from the EBI’s ENA FTP site for the three samples, using Run Accessions ERR903030, ERR895347 and ERR894724 for HG00514, HG00733 and NA19240, respectively. We ran the graph and linear mapping and genotyping pipelines exactly as for the simulation, and aggregated the comparison results across the three samples. We used BayesTyper to jointly genotype the 3 samples.

### GIAB Analysis

We obtained version 0.5 of the Genome in a Bottle (GIAB) SV VCF for the Ashkenazim son (HG002) and his parents from the NCBI FTP site. We obtained Illumina reads as described in Garrison et al.[15] and downsampled them to 50x coverage. We used these reads as input for `vg call` and the other SV genotyping pipelines described above (though with GRCh37 instead of GRCh38). For BayesTyper, we created the input variant set by combining the GIAB SVs with SNV and indels from the same study. Variants with reference allele or without a determined genotype for HG002 in the GIAB call set (10,569 out of 30,224) were considered “false positives” as a proxy measure for precision. These variants correspond to putative technical artifacts and parental calls not present in HG002. For the evaluation in high confidence regions, we used the Tier 1 high-confidence regions provided by the GIAB consortium in version 0.6.

### SMRT-SV v2 Comparison (CHMPD and SVPOP)

The SMRT-SV v2 Genotyper can only be used to genotype sequence-resolved SVs present on contigs with known SV breakpoints, such as those created by SMRT-SV v2, and therefore could not be run on the simulated, HGSVC, or GIAB call sets. The authors shared their training and evaluation set: a pseudodiploid sample constructed from combining the haploid CHM1 and CHM13 samples (CHMPD), and a negative control (NA19240). The high quality of the CHM assemblies makes this set an attractive alternative to using simulated reads. We used this two-sample pseudodiploid VCF along with the 30X read set to construct, map and genotype with vg, and also ran SMRT-SV v2 Genotyper with the “30x-4” model and min-call-depth 8 cutoff, and compared the two back to the original VCF.

In an effort to extend this comparison from the training data to a more realistic setting, we reran the three HGSVC samples against the SMRT-SV v2 discovery VCF (SVPOP, which contains 12 additional samples in addition to the three from HGSVC) published by Audano et al.[5] using vg and SMRT-SV v2 Genotyper. The discovery VCF does not contain genotypes. In consequence, we were unable to distinguish between heterozygous and homozygous genotypes, and instead considered only the presence or absence of a non-reference allele for each variant.

SMRT-SV v2 Genotyper produces explicit *no-call* predictions when the read coverage is too low to produce accurate genotypes. These no-calls are considered homozygous reference in the main accuracy evaluation. We also explored the performance of vg and SMRT-SV v2 Genotyper in different sets of regions (Figure S12 and Table S5):

1. Non-repeat regions, i.e. excluding segmental duplications and tandem repeats (using the respective tracks from the UCSC Genome Browser).
2. Repeat regions defined as segmental duplications and tandem repeats.
3. Regions where SMRT-SV v2 Genotyper could call variants.
4. Regions where SMRT-SV v2 Genotyper produced no-calls.

### Yeast graph analysis

For the analysis of graphs from *de novo* assemblies, we utilized publicly available PacBio-derived assemblies and Illumina short read sequencing datasets for 12 yeast strains from two related clades (Table 1) [28]. We constructed graphs from two different strain sets: For the *five strains set*, we selected five strains for graph construction (*S.c. SK1, S.c. YPS128, S.p. CBS432, S.p. UFRJ50816* and *S.c. S288C*). We randomly selected two strains from different subclades of each clade as well as the reference strain *S.c. S288C*. For the *all strains set* in contrast, we utilized all twelve strains for graph construction. We constructed two different types of genome graphs from the PacBio-derived assemblies of the five or twelve (depending on the strains set) selected strains. In this section, we describe the steps for the construction of both graphs and the genotyping of variants. More details and the precise commands used in our analyses can be found at github.com/vgteam/sv-genotyping-paper.

**Table 1:** 12 yeast strains from two related clades were used in our analysis. Five strains were selected to be included in the *five strains set* and all strains were included in the *all strains set*. Graphs were constructed from strains in the respective strain set while all eleven non-reference strains were used for genotyping.

| Strain | Clade | Included in *five strains set* | Included in *all strains set* |
| --- | --- | --- | --- |
| S288C | *S. cerevisiae* | ✓ | ✓ |
| SK1 | *S. cerevisiae* | ✓ | ✓ |
| YPS128 | *S. cerevisiae* | ✓ | ✓ |
| UWOPS034614 | *S. cerevisiae* |  | ✓ |
| Y12 | *S. cerevisiae* |  | ✓ |
| DBVPG6765 | *S. cerevisiae* |  | ✓ |
| DBVPG6044 | *S. cerevisiae* |  | ✓ |
| CBS432 | *S. paradoxus* | ✓ | ✓ |
| UFRJ50816 | *S. paradoxus* | ✓ | ✓ |
| N44 | *S. paradoxus* |  | ✓ |
| UWOPS919171 | *S. paradoxus* |  | ✓ |
| YPS138 | *S. paradoxus* |  | ✓ |

#### Construction of the *VCF graph*

We constructed the first graph (called the *VCF graph* throughout the paper) by adding variants onto a linear reference. This method requires one assembly to serve as a reference genome. The other assemblies must be converted to variant calls relative to this reference. The PacBio assembly of the S.c. S288C strain was chosen as the reference genome because this strain was used for the S. cerevisiae genome reference assembly. To obtain variants for the other assemblies, we combined three methods for SV detection from genome assemblies: Assemblytics [29] (commit df5361f), AsmVar (commit 5abd91a) [30] and paftools (version 2.14-r883) [31]. We constructed a union set of SVs detected by the three methods (using bedtools [43]), and combined variants with a reciprocal overlap of at least 50% to avoid duplication in the union set. We merged these union sets of variants for each of the other (non-reference) strains in the strain set, and we then applied another deduplication step to combine variants with a reciprocal overlap of at least 90%. We then used `vg construct` to build the *VCF graph* with the total set of variants and the linear reference genome.

#### Construction of the *cactus graph*

The second graph (called the *cactus graph* throughout the paper) was constructed from a whole genome alignment between the assemblies. First, the repeat-masked PacBio-assemblies of the strains in the strain set were aligned with our Cactus tool [27]. Cactus requires a phylogenetic tree of the strains which was estimated using Mash (version 2.1) [44] and PHYLIP (version 3.695) [45]. Subsequently, we converted the HAL format output Ale to a variation graph with hal2vg (https://github.com/ComparativeGenomicsToolkit/hal2vg).

#### Genotyping of SVs

Prior to genotyping, we mapped the Illumina short reads of all 12 yeast strains to both graphs using `vg map`. We measured the fractions of reads mapped with specific properties using `vg view` and the JSON processor `jq`. Then, we applied `toil-vg call` (commit be8b6da) to genotype variants, obtaining a separate genotype set for each of the 11 non-reference strains on both graphs and for each of the two strain sets (in total 11 x 2 x 2 = 44 genotype sets). From the genotype sets, we removed variants smaller than 50 bp and variants with missing or homozygous reference genotypes. To evaluate the Altered genotype sets, we generated a sample graph (i.e. a graph representation of the genotype set) for each genotype set using `vg construct` and `vg mod` on the reference assembly *S.c. S288C* and the genotype set. Subsequently, we mapped short reads from the respective strains to each sample graph using `vg map`. We mapped the short reads also to an empty sample graph that was generated using `vg construct` as a graph representation of the linear reference genome. In an effort to restrict our analysis to SV regions, we removed reads that mapped equally well (i.e. with identical mapping quality and percent identity) to all three graphs (the two sample graphs and the empty sample graph) from the analysis. These Altered out reads most likely stem from portions of the strains’ genomes that are identical to the reference strain *S.c. S288C*. We analyzed the remaining alignments of reads from SV regions with `vg view` and `jq`.

## Declarations

### Availability of data and material

The commands used to run the analyses presented in this study are available at github.com/vgteam/sv-genotyping-paper[46]. The datasets generated and/or analysed during the current study are also listed in this repository.

The scripts to generate the manuscript, including figures and tables, are available at github.com/jmonlong/manu-vgsv.

### Authors’ contributions

EG, AN, GH, JS, JE and ED implemented the read mapping and variant calling in the vg toolkit. GH, DH, JM, JAS and EG performed analysis on the different datasets. GH, DH, JM and BP designed the study. GH, DH and JM drafted the manuscript. All authors read, reviewed, and approved the Anal manuscript.

## Acknowledgements

We thank Peter Audano for sharing the CHMPD dataset and for his assistance with SMRT-SV v2.

## Authors’ information

These authors contributed equally: Glenn Hickey, David Heller, Jean Monlong.

## Supplementary Material

### Supplementary Tables

**Table S1:**
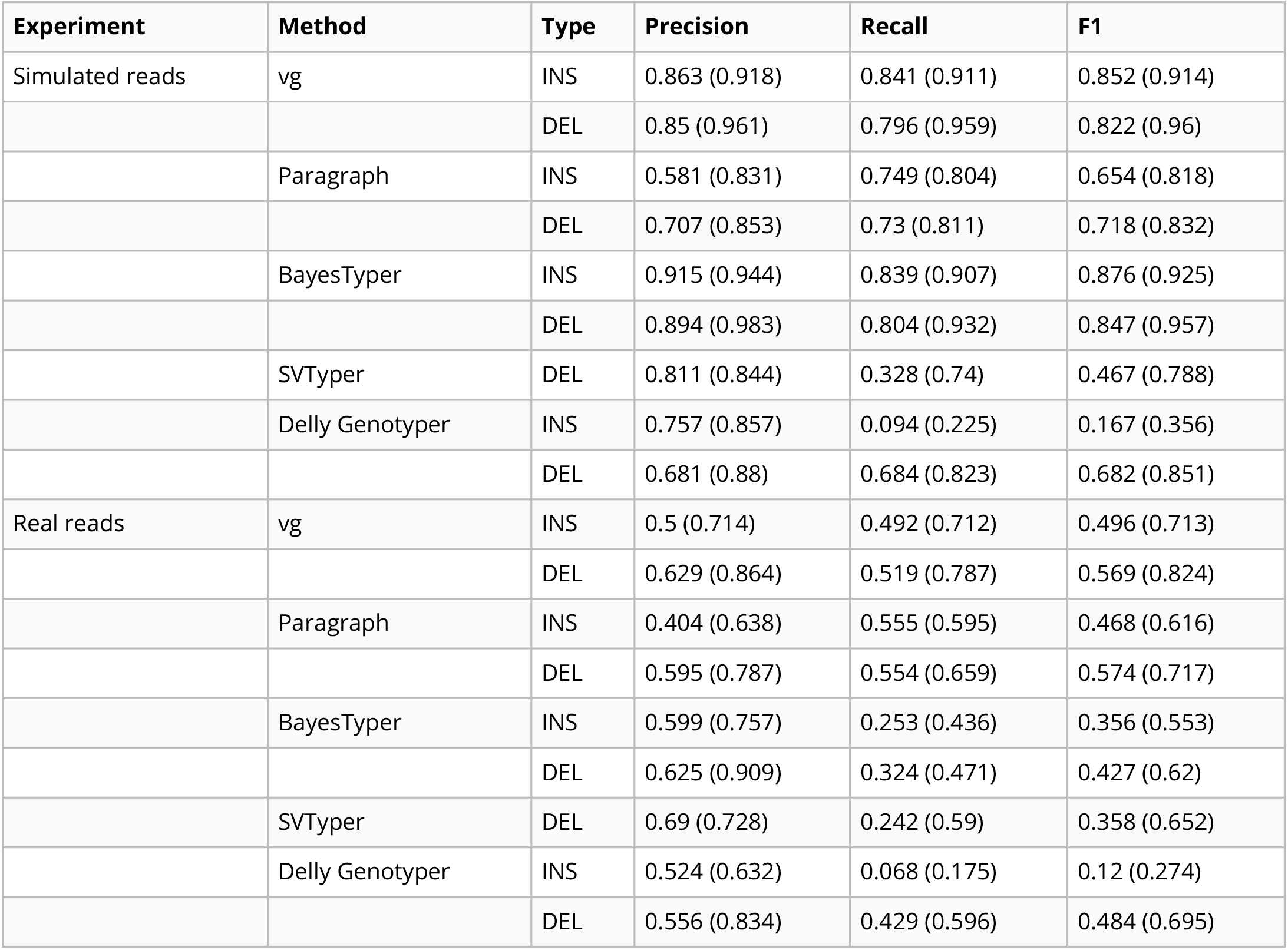
Genotyping evaluation on the HGSVC dataset. Precision, recall and F1 score for the call set with the best F1 score. The best F1 scores were achieved with no filtering in the vast majority of cases (see Figure S1 and S3). The numbers in parentheses corresponds to the results in non-repeat regions.

**Table S2:** Genotyping evaluation on the Genome in a Bottle dataset. Precision, recall and F1 score for the call set with the best F1 score. The best F1 scores were achieved with no filtering in the vast majority of cases (see Figure S7). The numbers in parentheses corresponds to the results in non-repeat regions.

| Method | Type | Precision | Recall | F1 |
| --- | --- | --- | --- | --- |
| vg | INS | 0.649 (0.776) | 0.618 (0.73) | 0.633 (0.752) |
|  | DEL | 0.696 (0.807) | 0.691 (0.795) | 0.694 (0.801) |
| Paragraph | INS | 0.699 (0.827) | 0.673 (0.768) | 0.686 (0.796) |
|  | DEL | 0.75 (0.9) | 0.726 (0.815) | 0.737 (0.855) |
| BayesTyper | INS | 0.777 (0.879) | 0.285 (0.379) | 0.417 (0.53) |
|  | DEL | 0.807 (0.884) | 0.514 (0.694) | 0.628 (0.778) |
| SVTyper | DEL | 0.743 (0.817) | 0.341 (0.496) | 0.467 (0.618) |
| Delly Genotyper | INS | 0.804 (0.888) | 0.178 (0.269) | 0.292 (0.413) |
|  | DEL | 0.721 (0.821) | 0.644 (0.766) | 0.68 (0.793) |

**Table S3:** Genotyping evaluation on the pseudo-diploid genome built from CHM cell lines in Audano et al.[5]. The numbers in parentheses corresponds to the results in non-repeat regions.

| Method | Type | Precision | Recall | F1 |
| --- | --- | --- | --- | --- |
| vg | INS | 0.783 (0.907) | 0.773 (0.895) | 0.778 (0.901) |
|  | DEL | 0.787 (0.962) | 0.635 (0.901) | 0.703 (0.93) |
| SMRT-SV v2 Genotyper | INS | 0.819 (0.934) | 0.582 (0.712) | 0.681 (0.808) |
|  | DEL | 0.848 (0.973) | 0.63 (0.839) | 0.723 (0.901) |

**Table S4:** Calling evaluation on the SVPOP dataset. Combined results for the HG00514, HG00733 and NA19240 individuals, 3 of the 15 individuals used to generate the high-quality SV catalog in Audano et al.[5].

| Method | Region | Type | TP | FP | FN | Precision | Recall | F1 |
| --- | --- | --- | --- | --- | --- | --- | --- | --- |
| vg | all | INS | 23430 | 18414 | 18181 | 0.564 | 0.563 | 0.564 |
|  |  | DEL | 14717 | 7033 | 15254 | 0.677 | 0.491 | 0.569 |
|  |  | INV | 41 | 16 | 159 | 0.719 | 0.205 | 0.319 |
|  | non-repeat | INS | 8078 | 3303 | 1761 | 0.709 | 0.821 | 0.761 |
|  |  | DEL | 6585 | 1033 | 1040 | 0.862 | 0.864 | 0.863 |
|  |  | INV | 37 | 15 | 90 | 0.712 | 0.291 | 0.413 |
| Paragraph | all | INS | 24342 | 25618 | 17269 | 0.493 | 0.585 | 0.535 |
|  |  | DEL | 16986 | 13376 | 12985 | 0.571 | 0.567 | 0.569 |
|  |  | INV | 47 | 24 | 153 | 0.662 | 0.235 | 0.347 |
|  | non-repeat | INS | 7843 | 3270 | 1996 | 0.706 | 0.797 | 0.749 |
|  |  | DEL | 6523 | 1000 | 1102 | 0.866 | 0.856 | 0.860 |
|  |  | INV | 39 | 12 | 88 | 0.765 | 0.307 | 0.438 |
| SMRT-SV v2 Genotyper | all | INS | 16297 | 26006 | 25314 | 0.397 | 0.392 | 0.394 |
|  |  | DEL | 11797 | 10054 | 18174 | 0.544 | 0.394 | 0.457 |
|  | non-repeat | INS | 4475 | 4645 | 5364 | 0.493 | 0.455 | 0.473 |
|  |  | DEL | 4986 | 1322 | 2639 | 0.788 | 0.654 | 0.715 |

**Table S5:** Calling evaluation on the SVPOP dataset in different sets of regions for the HG5014 individual.

| Method | Region | Type | TP | FP | FN | Precision | Recall | F1 |
| --- | --- | --- | --- | --- | --- | --- | --- | --- |
| vg | all | INS | 7764 | 6109 | 6270 | 0.567 | 0.553 | 0.560 |
|  |  | DEL | 4841 | 2260 | 5066 | 0.684 | 0.489 | 0.570 |
|  |  | INV | 16 | 6 | 49 | 0.727 | 0.246 | 0.368 |
|  | repeat | INS | 5091 | 5150 | 5766 | 0.507 | 0.469 | 0.487 |
|  |  | DEL | 2684 | 1922 | 4648 | 0.590 | 0.366 | 0.452 |
|  |  | INV | 1 | 0 | 9 | 1.000 | 0.100 | 0.182 |
|  | non-repeat | INS | 2662 | 979 | 521 | 0.732 | 0.836 | 0.781 |
|  |  | DEL | 2085 | 322 | 388 | 0.865 | 0.843 | 0.854 |
|  |  | INV | 14 | 6 | 26 | 0.700 | 0.350 | 0.467 |
|  | called in SMRT-SV v2 Genotyper | INS | 3682 | 4752 | 1836 | 0.444 | 0.667 | 0.534 |
|  |  | DEL | 2769 | 1779 | 1356 | 0.609 | 0.671 | 0.639 |
|  |  | INV | 16 | 6 | 49 | 0.727 | 0.246 | 0.368 |
|  | not called in SMRT-SV v2 Genotyper | INS | 3867 | 291 | 4649 | 0.931 | 0.454 | 0.610 |
|  |  | DEL | 1976 | 102 | 3797 | 0.952 | 0.342 | 0.503 |
| SMRT-SV v2 Genotyper | all | INS | 5254 | 8562 | 8780 | 0.394 | 0.374 | 0.384 |
|  |  | DEL | 3743 | 3367 | 6164 | 0.535 | 0.378 | 0.443 |
|  | repeat | INS | 3858 | 7119 | 6999 | 0.368 | 0.355 | 0.362 |
|  |  | DEL | 2141 | 2906 | 5191 | 0.438 | 0.292 | 0.350 |
|  | non-repeat | INS | 1394 | 1464 | 1789 | 0.493 | 0.438 | 0.464 |
|  |  | DEL | 1550 | 443 | 923 | 0.778 | 0.627 | 0.694 |
|  | called in SMRT-SV v2 Genotyper | INS | 4360 | 5619 | 1158 | 0.445 | 0.790 | 0.570 |
|  |  | DEL | 3272 | 2554 | 853 | 0.568 | 0.793 | 0.662 |
|  | not called in SMRT-SV v2 Genotyper | INS | 111 | 101 | 8405 | 0.549 | 0.013 | 0.025 |
|  |  | DEL | 211 | 50 | 5562 | 0.792 | 0.036 | 0.070 |

**Table S6:** Breakpoint fine-tuning using graph augmentation from the read alignment. For deletions and inversions, either one or both breakpoints were shifted to introduce errors in the input VCF. For insertions, the insertion location and sequence contained errors. In all cases, the errors affected 1-10 bp.

| SV type | Error type | Breakpoint | Variant | Proportion | Mean size (bp) | Mean error (bp) |
| --- | --- | --- | --- | --- | --- | --- |
| DEL | one end | incorrect | 220 | 0.219 | 422.655 | 6.095 |
|  |  | fine-tuned | 784 | 0.781 | 670.518 | 5.430 |
|  | both ends | incorrect | 811 | 0.814 | 826.070 | 6.275 |
|  |  | fine-tuned | 185 | 0.186 | 586.676 | 2.232 |
| INS | location/seq | incorrect | 123 | 0.062 | 428.724 | 6.667 |
|  |  | fine-tuned | 1877 | 0.938 | 440.043 | 6.439 |
| INV | one end | incorrect | 868 | 0.835 | 762.673 | 5.161 |
|  |  | fine-tuned | 172 | 0.165 | 130.244 | 5.884 |
|  | both ends | incorrect | 950 | 0.992 | 556.274 | 5.624 |
|  |  | fine-tuned | 8 | 0.008 | 200.000 | 1.375 |

## Supplementary Figures

**Figure S1:**
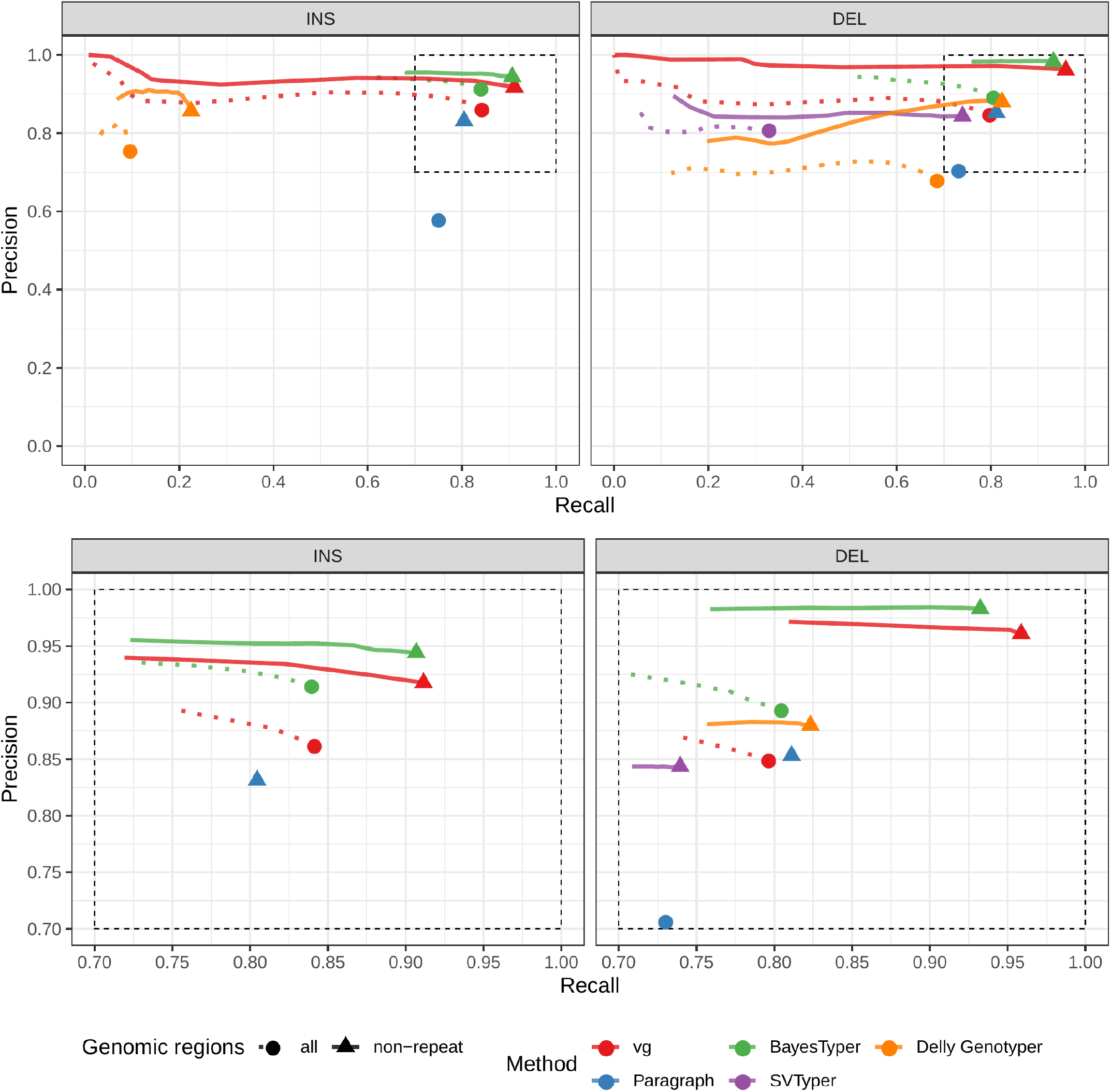
Genotyping evaluation on the HGSVC dataset using simulated reads. Reads were simulated from the HG00514 individual. The bottom panel zooms on the part highlighted by a dotted rectangle.

**Figure S2:**
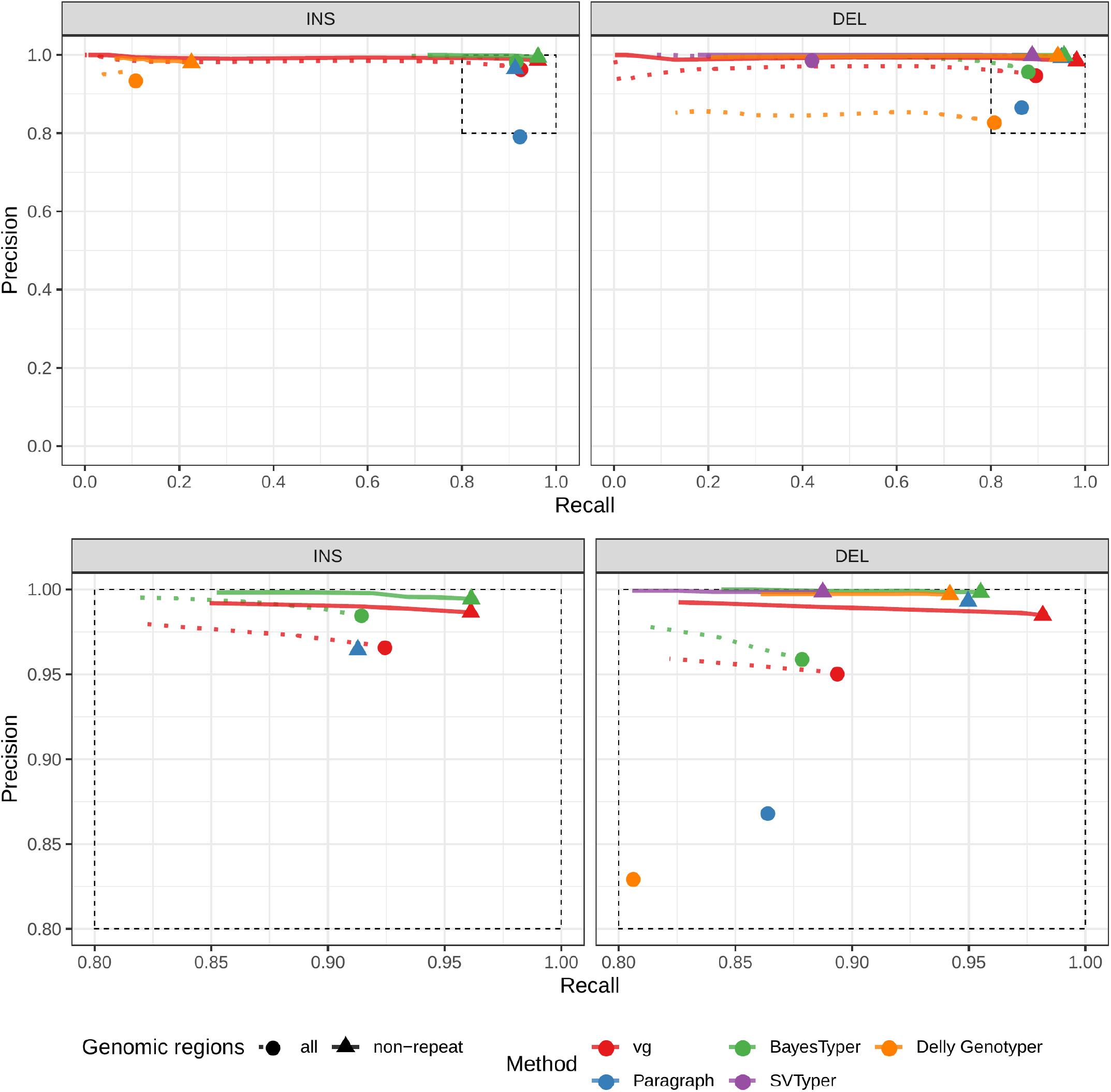
Calling evaluation on the HGSVC dataset using simulated reads. Reads were simulated from the HG00514 individual. The bottom panel zooms on the part highlighted by a dotted rectangle.

**Figure S3:**
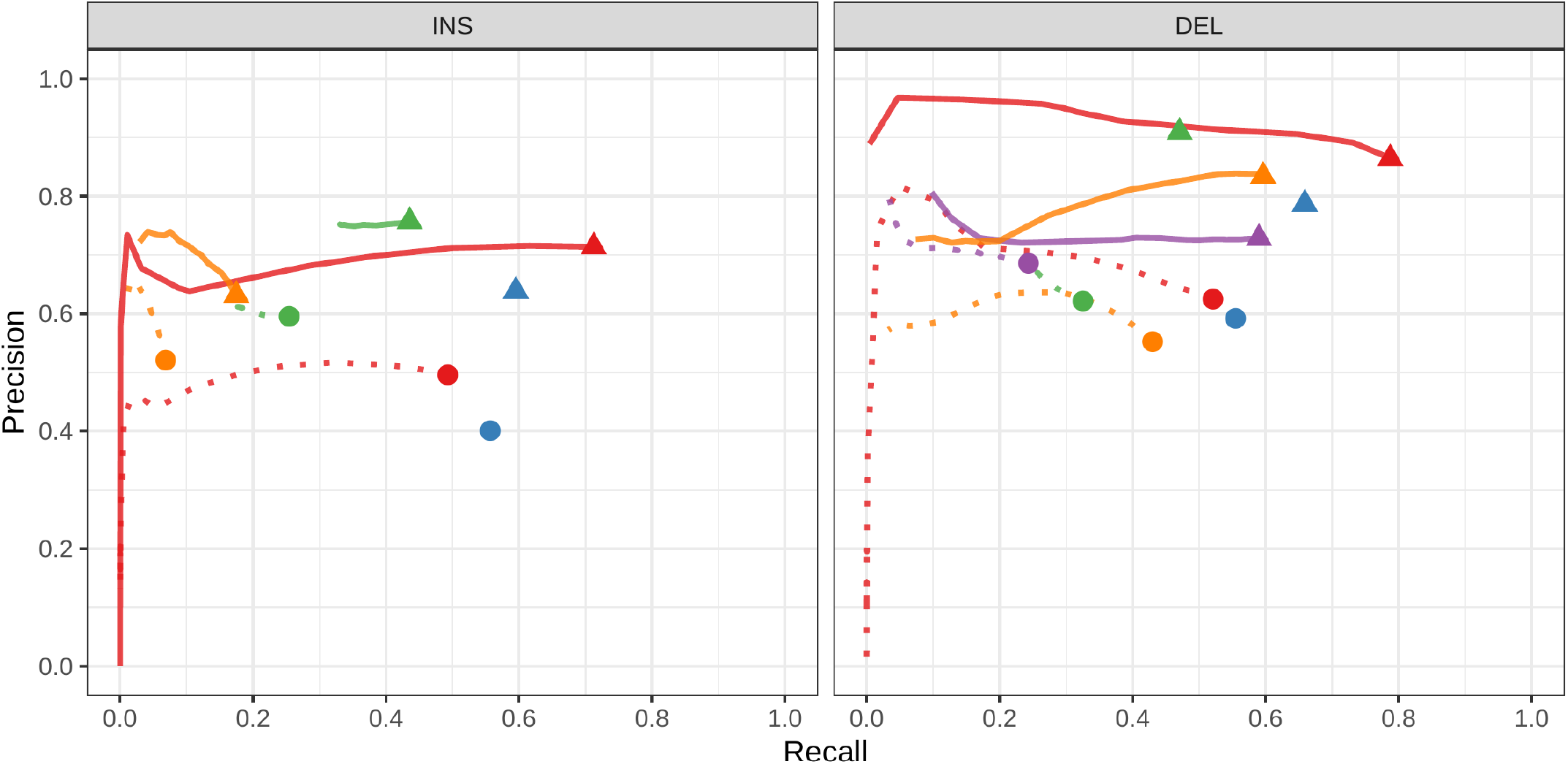
Genotyping evaluation on the HGSVC dataset using real reads. Combined results across the HG00514, HG00733 and NA19240.

**Figure S4:**
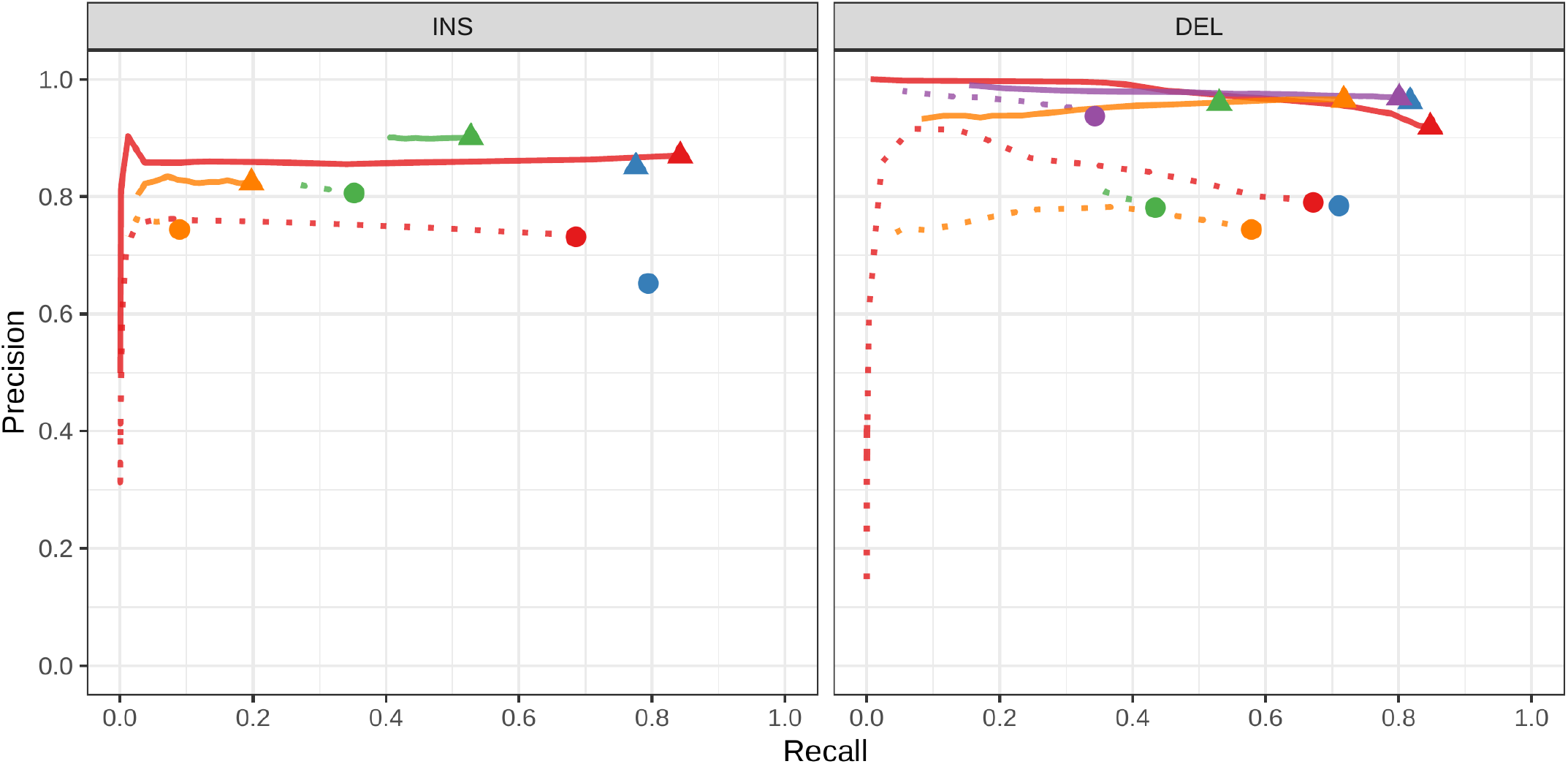
Calling evaluation on the HGSVC dataset using real reads. Combined results across the HG00514, HG00733 and NA19240.

**Figure S5:**
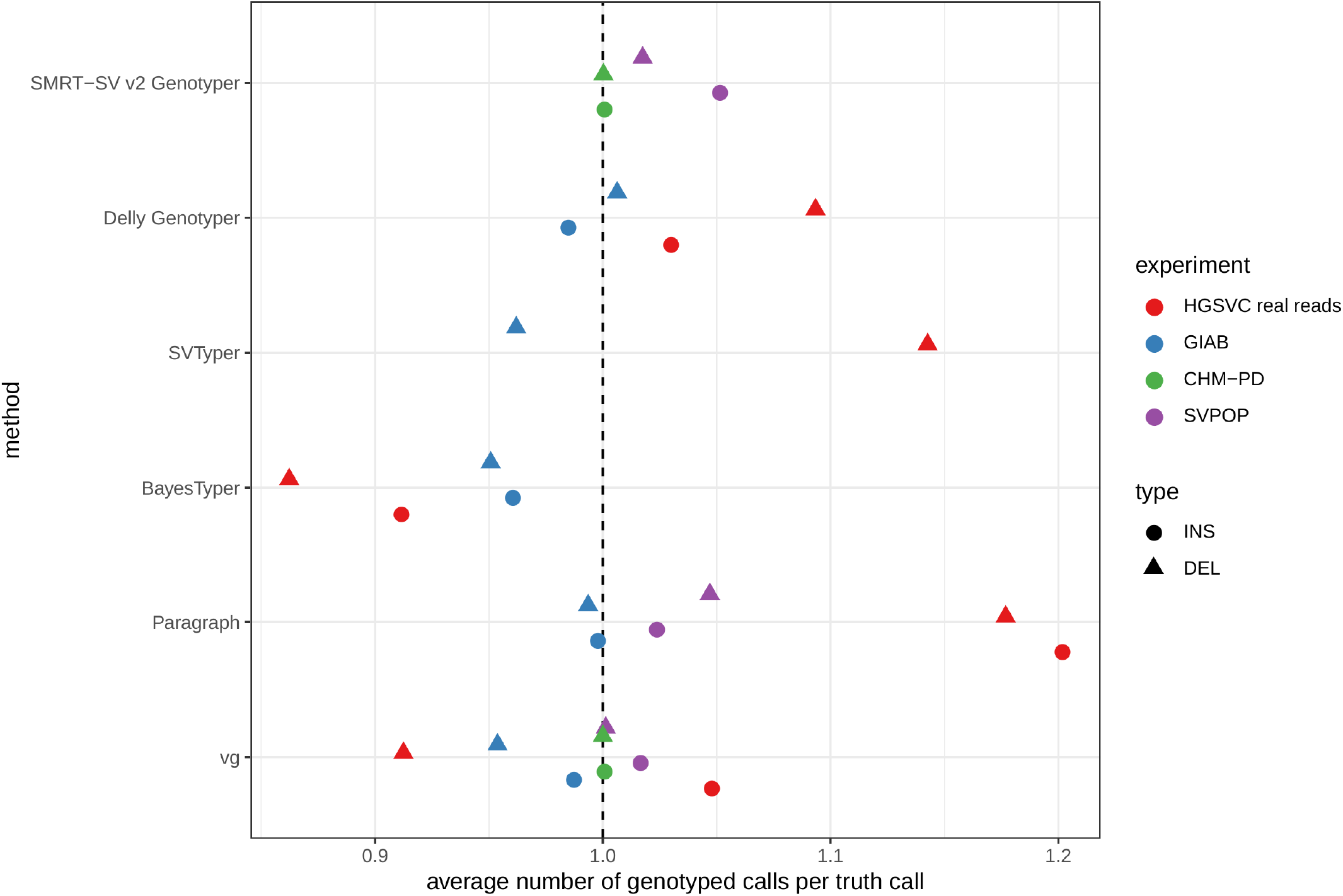
Average number of genotyped variants overlapping one variant from the truth set. To evaluate the genotyping performance, each genotyped variant is matched to variants in the truth set. A same variant can match to several variant in the other set because of variant fragmentation or when the truth set contains potentially duplicated SVs. This x-axis shows the average number of genotyped variants that were matched per truth-set variant. For example, a value higher than 1 means that variants in the truth were often matched to multiple genotyped variants (“over-genotyping”).

**Figure S6:**
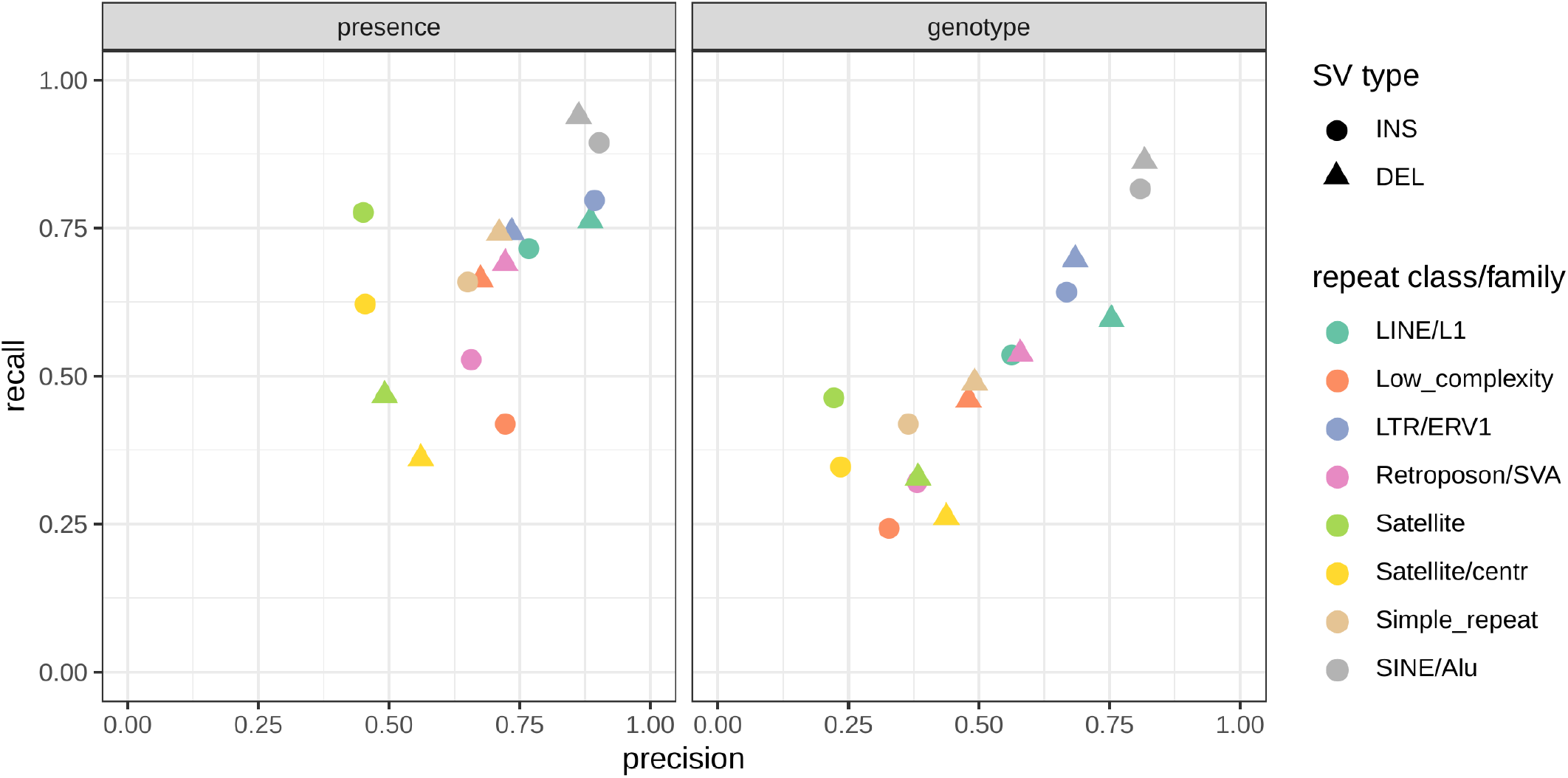
Evaluation across different repeat profiles. The deleted/inserted sequence was annotated with RepeatMasker (color). The precision and recall was recomputed on each of the most frequent repeat families.

**Figure S7:**
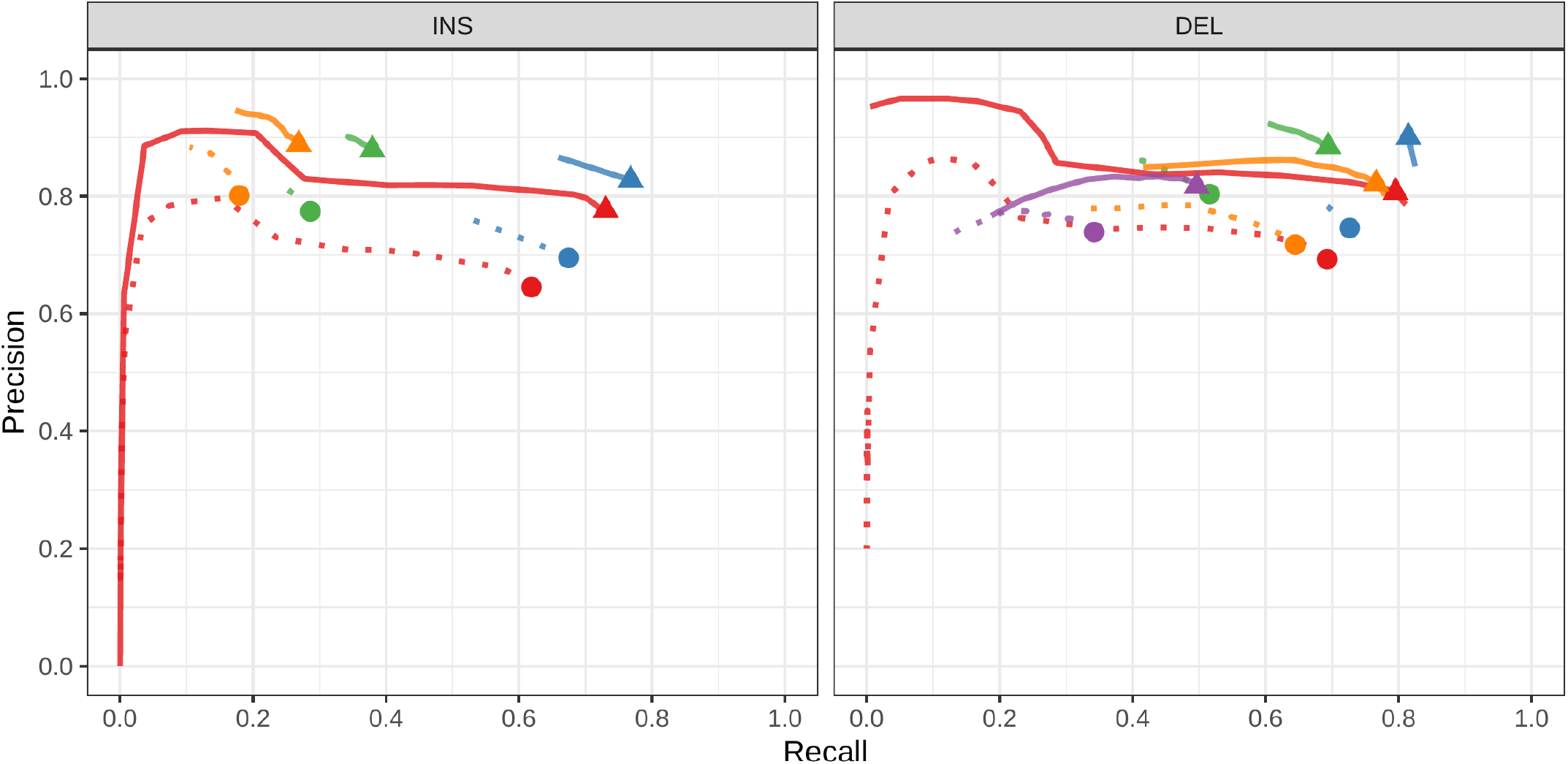
Genotyping evaluation on the Genome in a Bottle dataset. Predicted genotypes on HG002 were compared to the high-quality SVs from this same individual.

**Figure S8:**
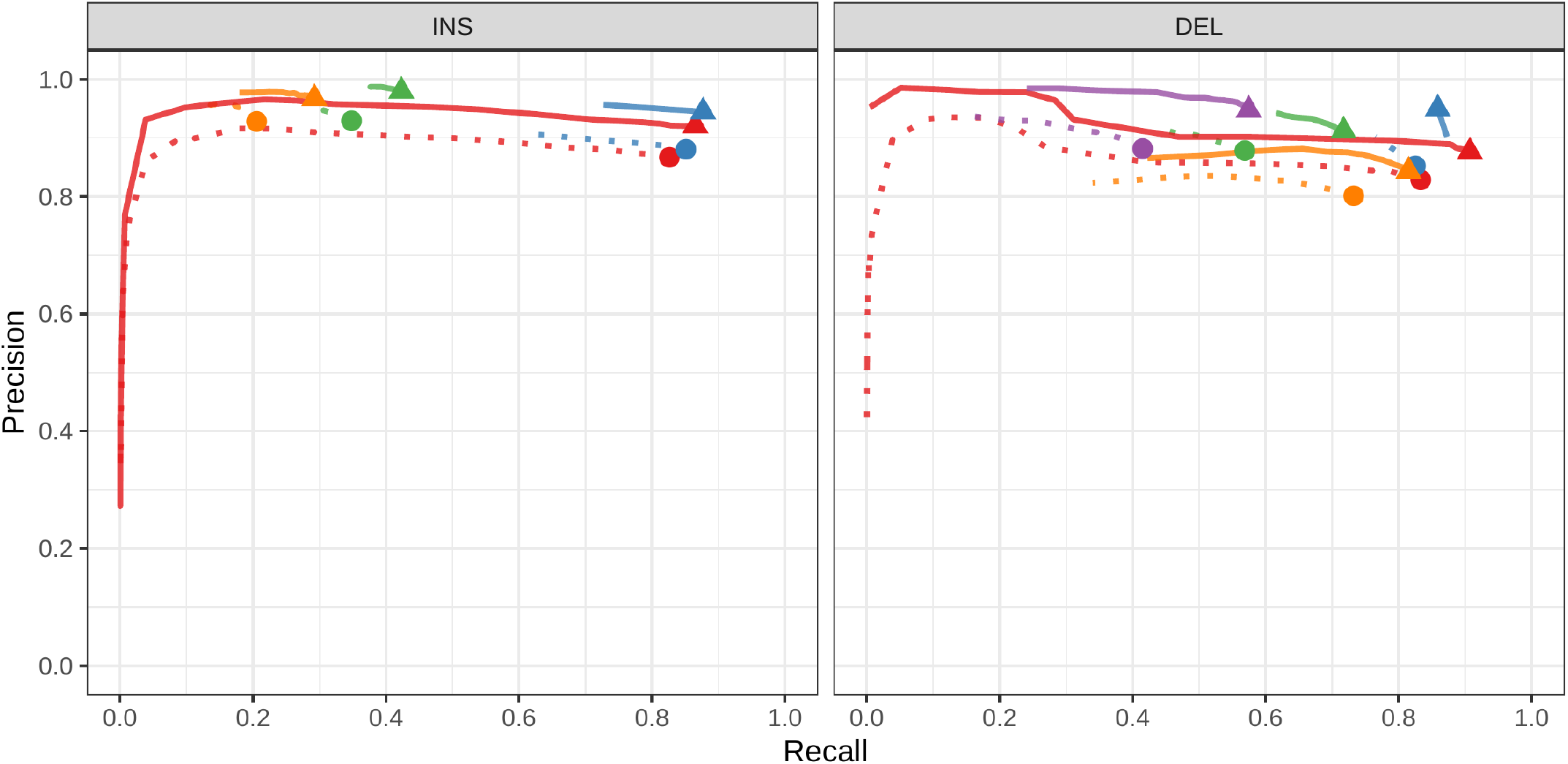
Calling evaluation on the Genome in a Bottle dataset. Calls on HG002 were compared to the high-quality SVs from this same individual.

**Figure S9:**
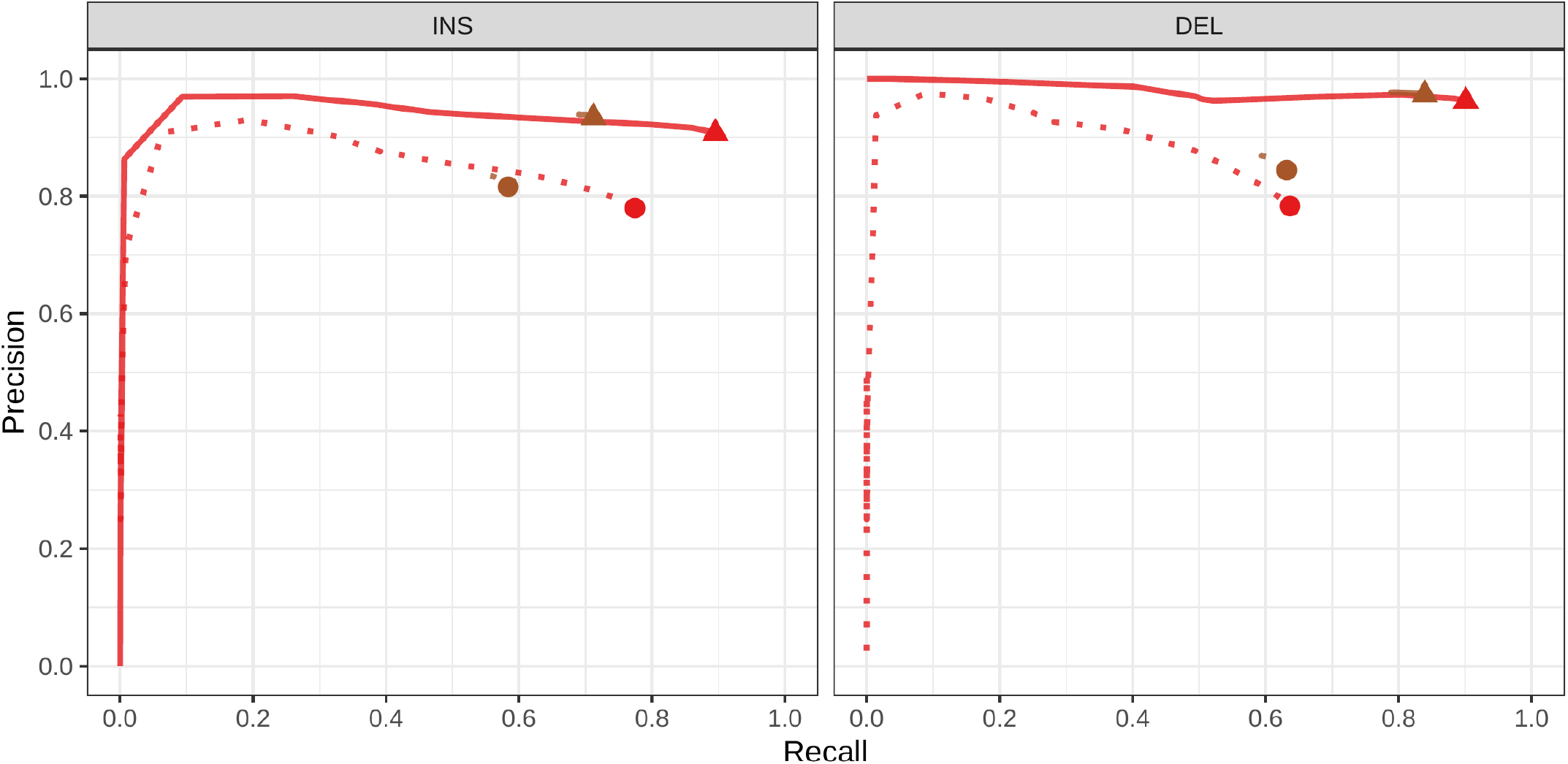
Genotyping evaluation on the CHM pseudo-diploid dataset. The pseudo-diploid genome was built from CHM cell lines and used to train SMRT-SV v2 Genotyper in Audano et al.[5] The bottom panel zooms on the part highlighted by a dotted rectangle.

**Figure S10:**
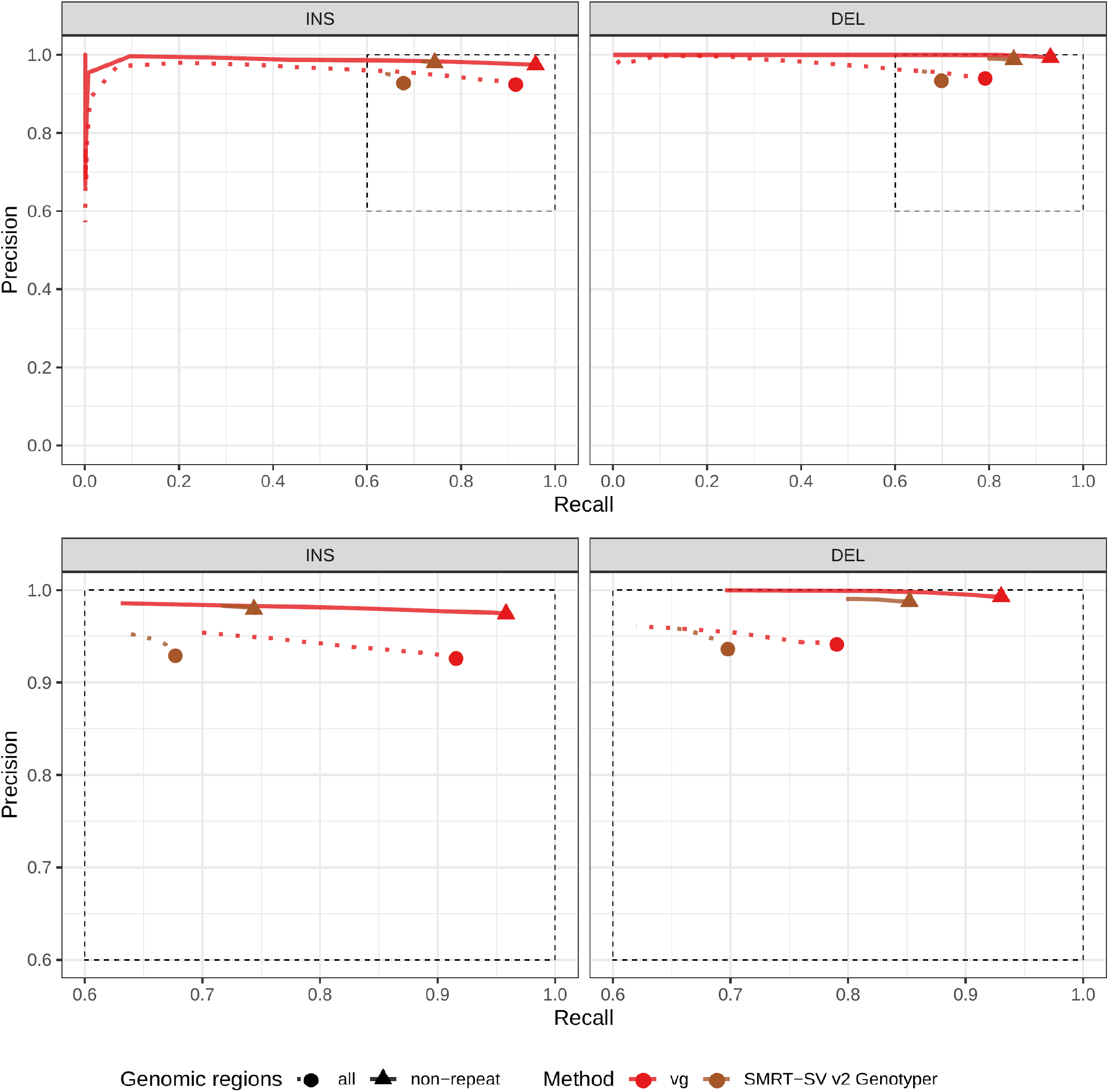
Calling evaluation on the CHM pseudo-diploid dataset. The pseudo-diploid genome was built from CHM cell lines and used to train SMRT-SV v2 Genotyper in Audano et al.[5]

**Figure S11:**
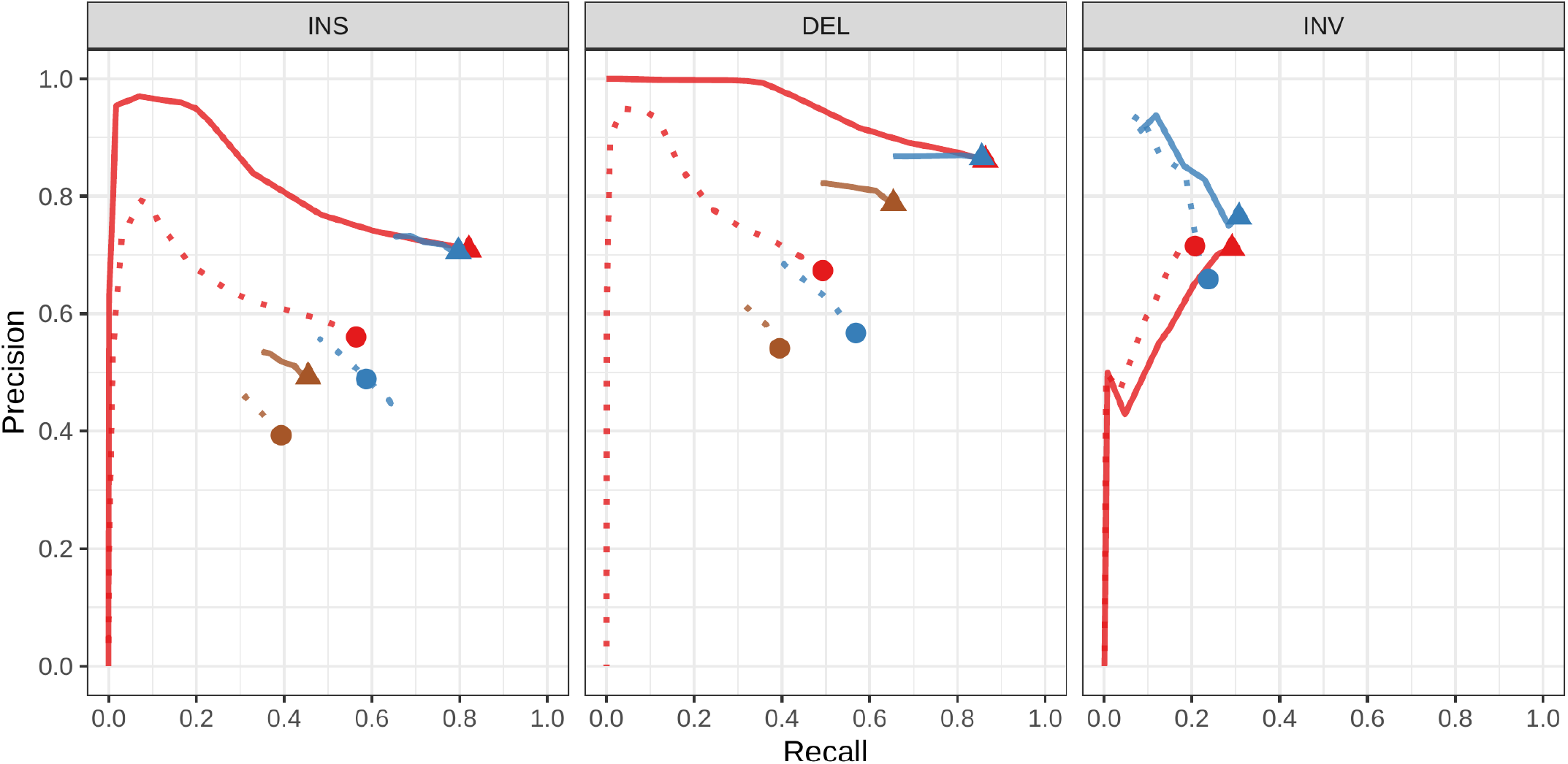
Calling evaluation on the SVPOP dataset. Combined results across the HG00514, HG00733 and NA19240.

**Figure S12:**
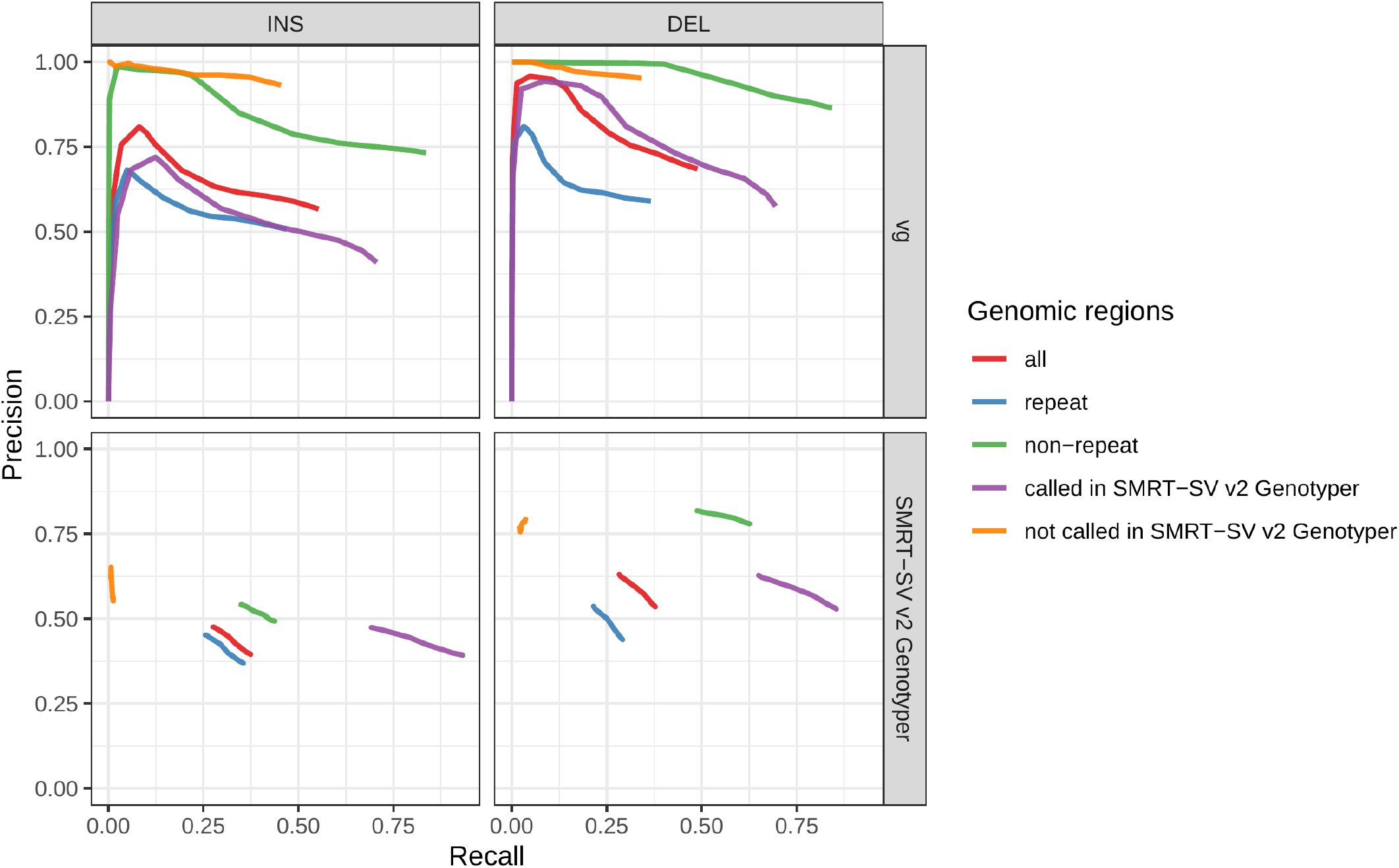
Evaluation across different sets of regions in HG00514 (SVPOP dataset). Calling evaluation.

**Figure S13:**
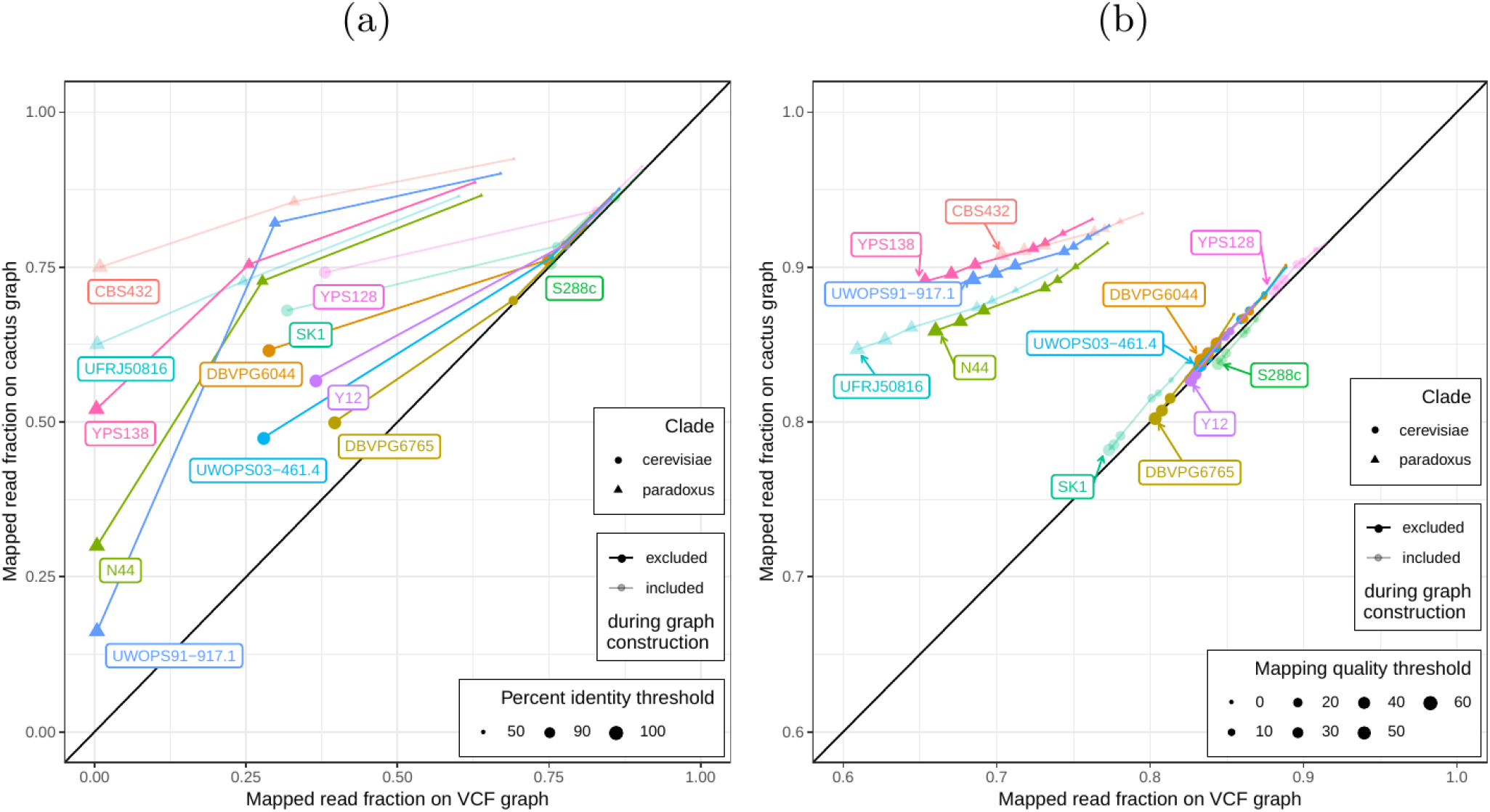
Mapping comparison on graphs of the *five strains set*. Short reads from all 12 yeast strains were aligned to both graphs. The fraction of reads mapped to the cactus graph (y-axis) and the VCF graph (x-axis) are compared. a) Stratified by percent identity threshold. b) Stratified by mapping quality threshold. Colors and shapes represent the 12 strains and two clades, respectively. Transparency indicates whether the strain was included or excluded in the graphs.

**Figure S14:**
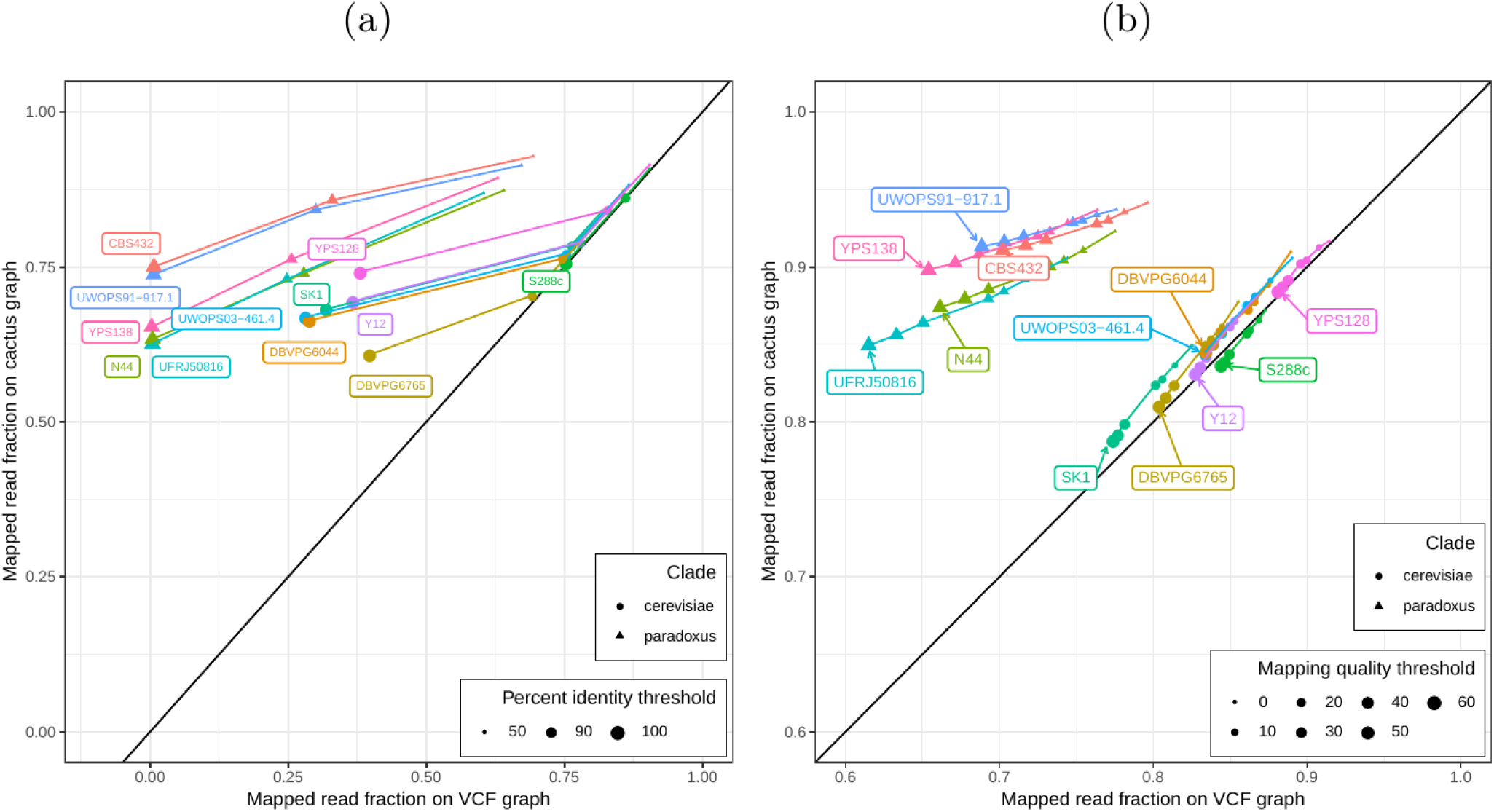
Mapping comparison on graphs of the *all strains set*. Short reads from all 12 yeast strains were aligned to both graphs. The fraction of reads mapped to the *cactus graph* (y-axis) and the *VCF graph* (x-axis) are compared. a) Stratified by percent identity threshold. b) Stratified by mapping quality threshold. Colors and shapes represent the 12 strains and two clades, respectively.

**Figure S15:**
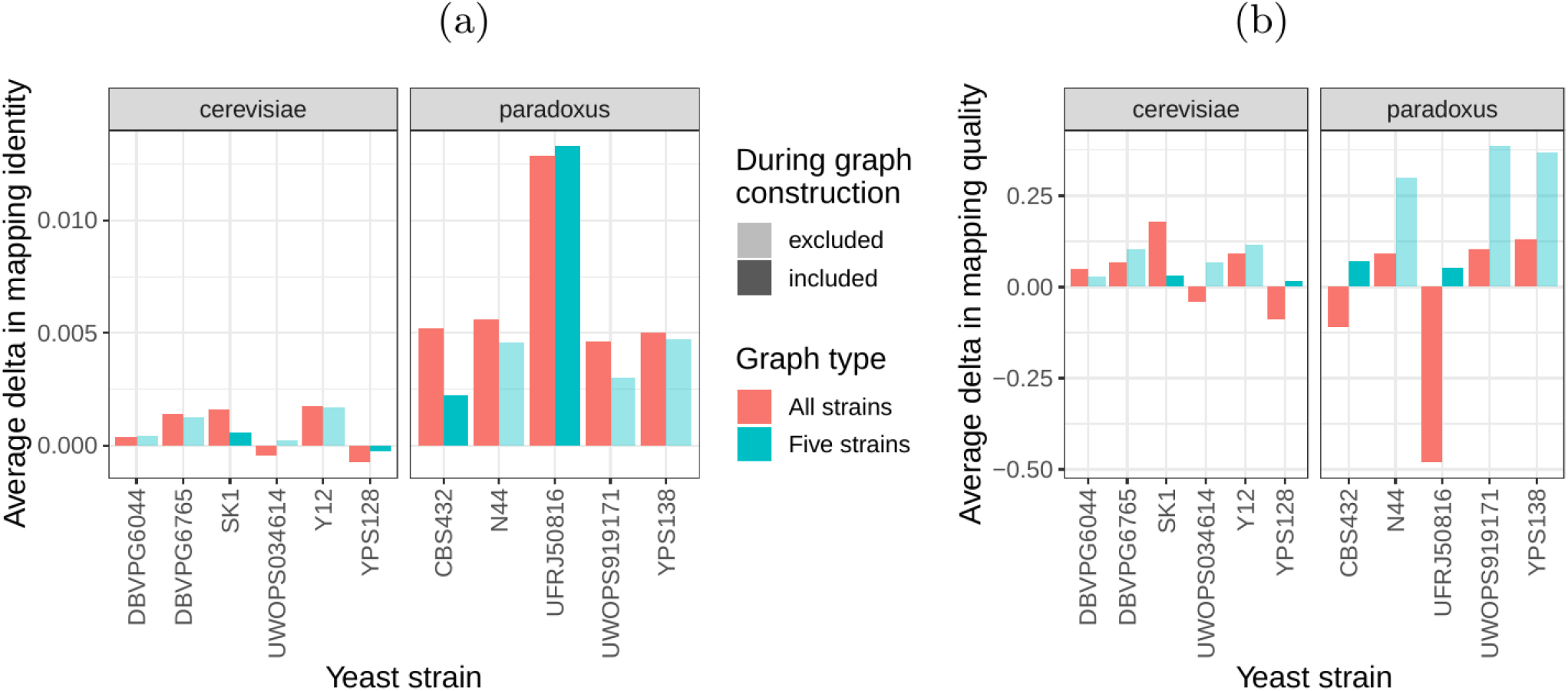
SV genotyping comparison using all reads. Short reads from all 11 non-reference yeast strains were used to genotype SVs contained in the *cactus graph* and the *VCF graph*. Subsequently, sample graphs were generated from the resulting SV callsets. The short reads were aligned to the sample graphs and the quality of all alignments was used to ascertain SV genotyping performance. More accurate genotypes should result in sample graphs that have mappings with high identity and confidence for a greater proportion of the reads. a) Average delta in mapping identity of all short reads aligned to the sample graphs derived from *cactus graph* and *VCF graph*. b) Average delta in mapping quality of all short reads aligned to the sample graphs derived from *cactus graph* and *VCF graph*. Positive values denote an improvement of the *cactus graph* over the *VCF graph*. Colors represent the two strain sets and transparency indicates whether the respective strain was part of the *five strains set*.

**Figure S16:**
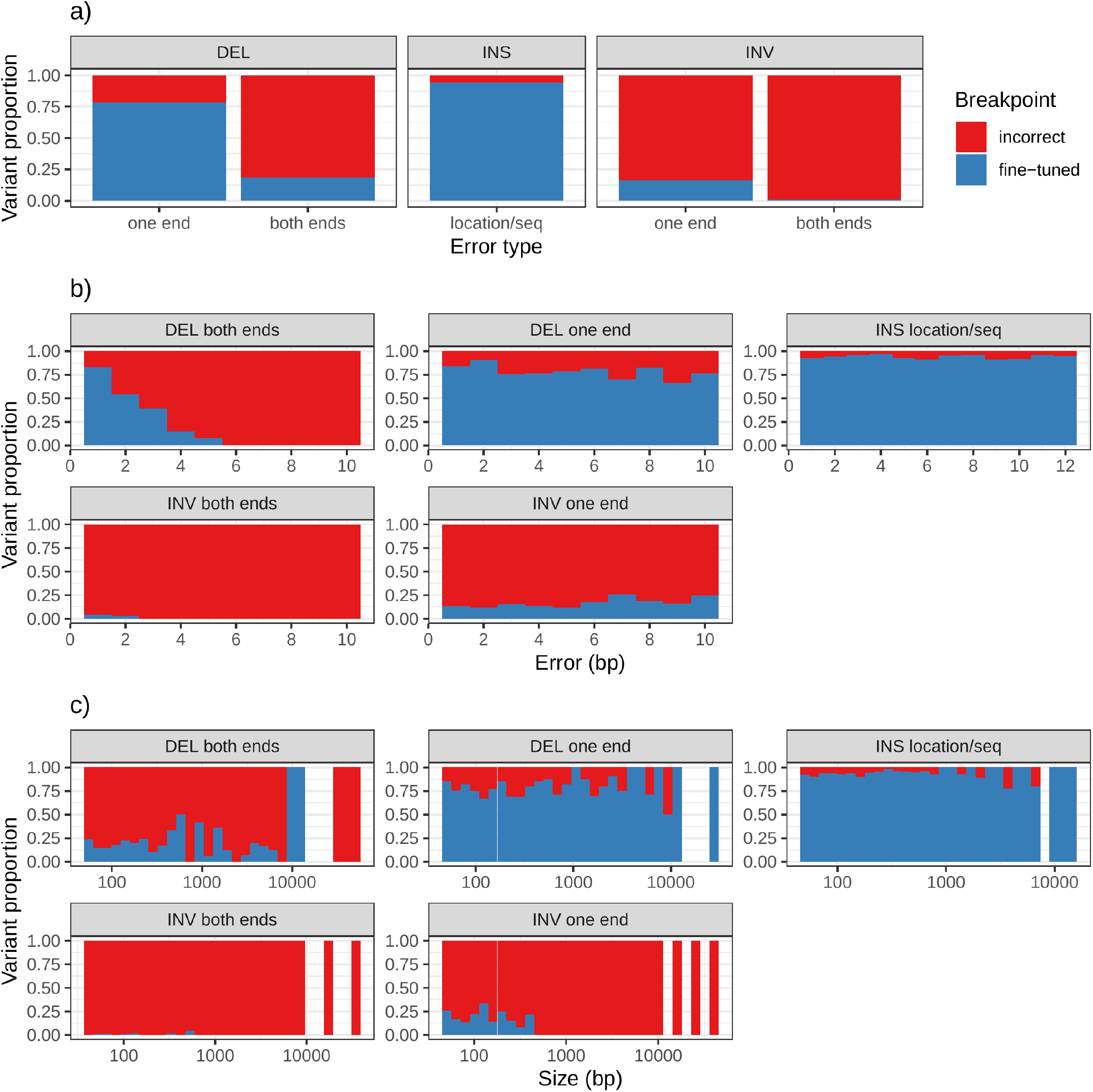
Breakpoint fine-tuning using augmentation through “vg call”. For deletions and inversions, either one or both breakpoints were shifted to introduce errors in the input VCF. For insertions, the insertion location and sequence contained errors. a) Proportion of variant for which breakpoints could be fine-tuned. b) Distribution of the amount of errors that could be corrected or not. c) Distribution of the size of the variants whose breakpoints could be fine-tuned or not.

**Figure S17:**
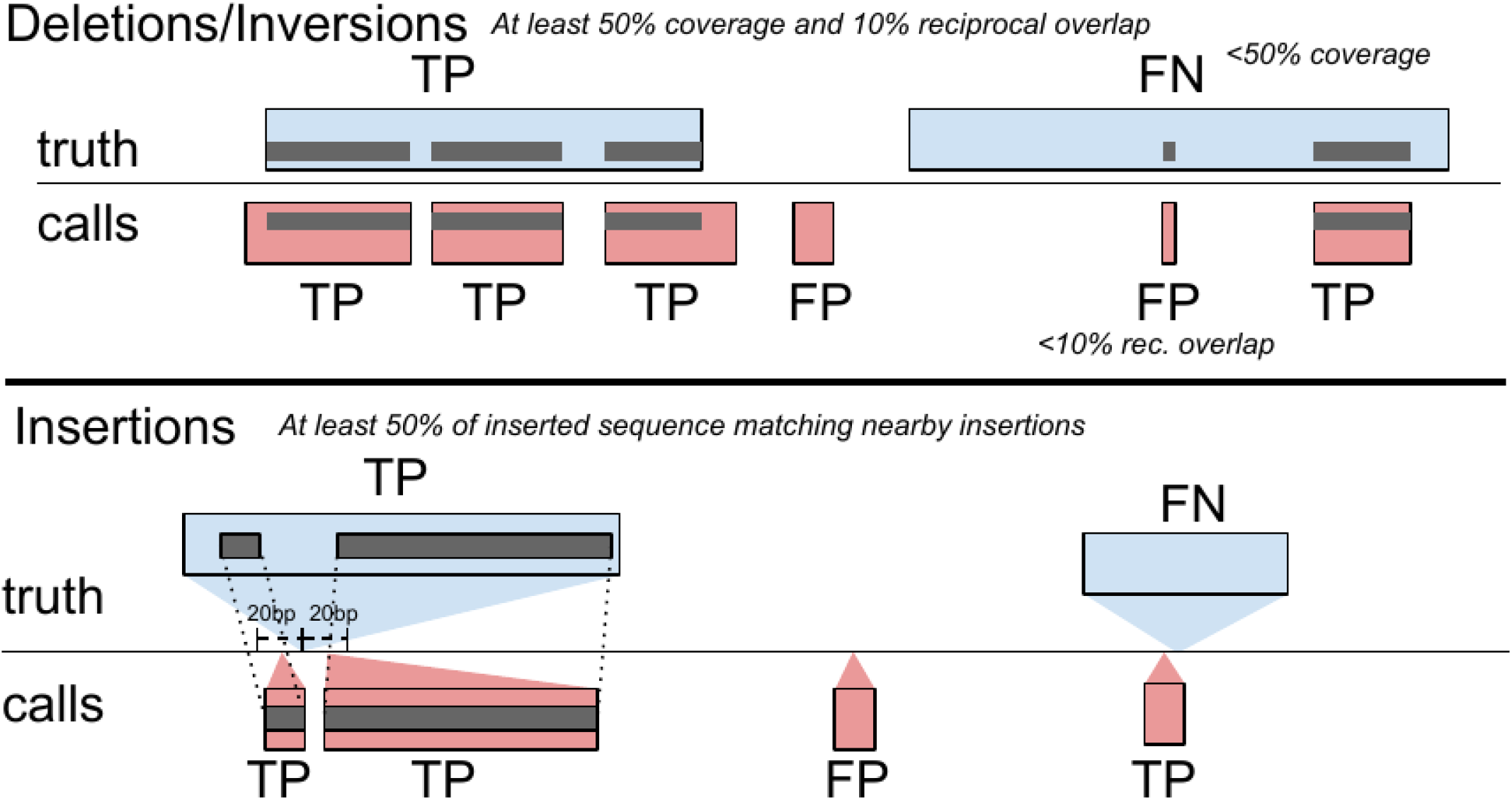
Overview of the SV evaluation by the *sveval* package. For deletions and inversions, we compute the proportion of a variant that is covered by variants in the other set, considering only variants overlapping with at least 10% reciprocal overlap. A variant is considered true positive if this coverage proportion is higher than 50% and falsepositive or false-negative otherwise. A similar approach is used for insertions, although they are first clustered into pairs located less than 20 bp from each other. Then their inserted sequences are aligned to derive the coverage statistics. The SV evaluation approach is described in more detail in the Methods.

**Figure S18:**
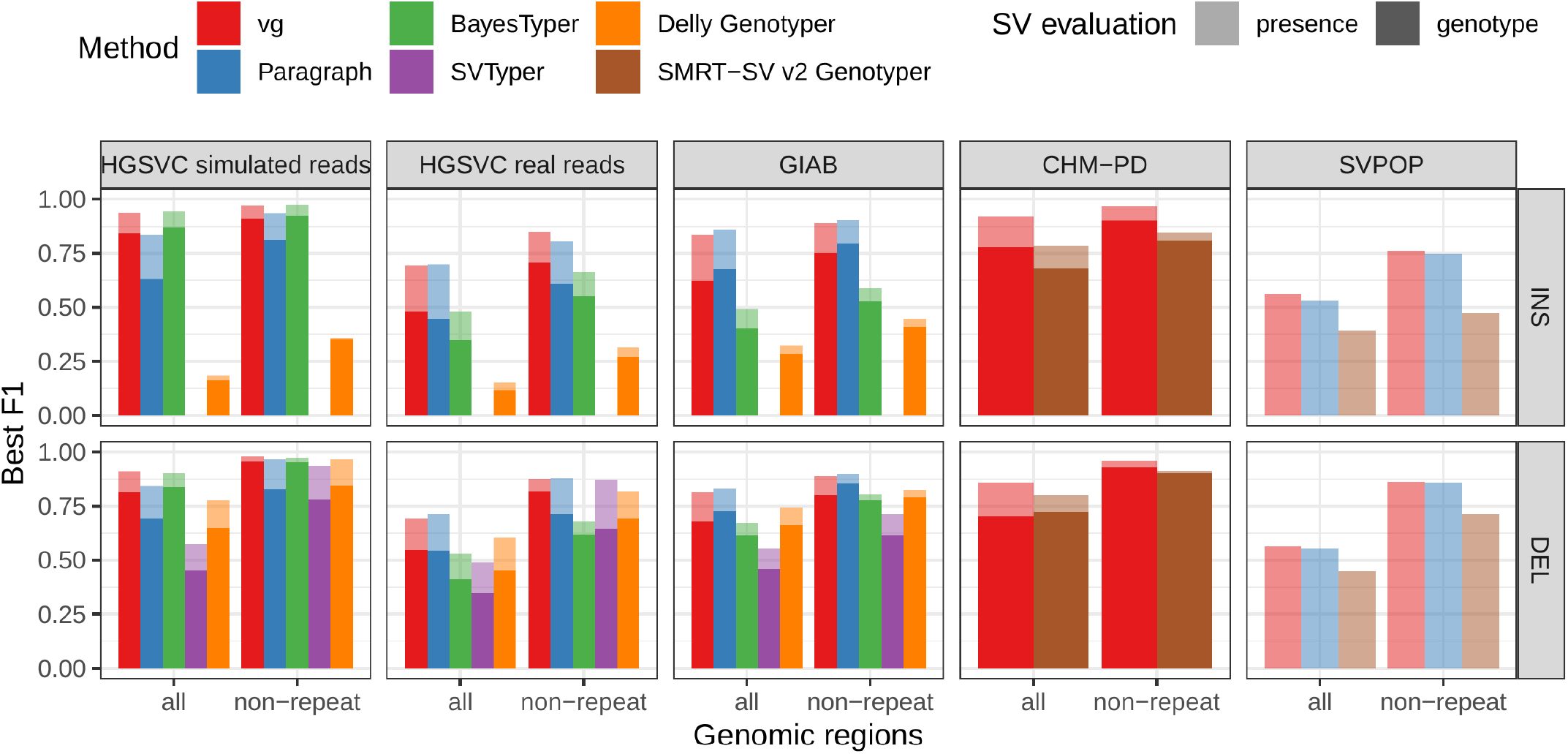
Benchmark summary when using a more stringent matching criterion. At least 90% coverage was necessary to consider a variant matched, instead of the 50% minimum coverage used in other figures.

## Supplementary Information

### Variation graph and structural variation

A variation graph encodes DNA sequence in its nodes. Such graphs are bidirected, in that we distinguish between edges incident on the starts of nodes from those incident on their ends. A path in such a graph is an ordered list of nodes where each is associated with an orientation. If a path walks from, for example, node A in the forward orientation to node B in the reverse orientation, then an edge must exist from the end of node A to the end of node B. Concatenating the sequences on each node in the path, taking the reverse complement when the node is visited in reverse orientation, produces a DNA sequence. Accordingly, variation graphs are constructed so as to encode haplotype sequences as walks through the graph. Variation between sequences shows up as bubbles in the graph [24].

### Breakpoint fine-tuning

In addition to genotyping, vg can use an augmentation step to modify the graph based on the read alignment and discover novel variants. On the simulated SVs from Figure 1b, this approach was able to correct many of the 1-10 bp breakpoint errors that were added to the input VCF. The breakpoints were accurately fine-tuned for 93.8% of the insertions (Figure S16a and Table S6). For deletions, 78.1% of the variants were corrected when only one breakpoint had an error. In situations where both breakpoints of the deletions were incorrect, only 18.6% were corrected through graph augmentation, and only when the amount of error was small (Figure S16b). The breakpoints of less than 20% of the inversions could be corrected. Across all SV types, the size of the variant didn’t affect the ability to fine-tune the breakpoints through graph augmentation (Figure S16c).

### Mappability comparison between yeast graphs

In order to elucidate whether the *cactus graph* represents the sequence diversity among the yeast strains better than the *VCF graph*, we mapped Illumina short reads to both graphs using `vg map`. Generally, more reads mapped to the *cactus graph* with high identity (Figures S13a and S14a) and high mapping quality (Figures S13b and S14b) than to the *VCF graph*. The *VCF graph* exhibited higher mappability only on the reference strain *S.c. S288C* with a marginal difference. The benefit of using the *cactus graph* is largest for strains in the *S. paradoxus* clade and smaller for strains in the *S. cerevisiae* clade. We found that the genetic distance to the reference strain (as estimated using Mash v2.1 [44]) correlated with the increase in confidently mapped reads (mapping quality >= 60) between the *cactus graph* and the *VCF graph* (Spearman’s rank correlation, p-value=3.993e-06). These results suggest that the improvement in mappability is not driven by the higher sequence content in the *cactus graph* alone (16.8 / 15.4 Mb in the *cactus graph* compared to 12.6 / 12.4 Mb in the *VCF graph* for the *all strains set* and the *five strains set*, respectively). Instead, an explanation could be the construction of the *VCF graph* from a comprehensive but still limited list of variants and the lack of SNPs and small Indels in this list. Consequently, substantially fewer reads mapped to the *VCF graph* with perfect identity (Figures S13a and S14a, percent identity threshold = 100%) than to the *cactus graph*. The *cactus graph* has the advantage of implicitly incorporating variants of all types and sizes from the *de novo* assemblies. As a consequence, the *cactus graph* captures the genetic makeup of each strain more comprehensively and enables more reads to be mapped.

Interestingly, our measurements for the *five strains set* showed only small differences between the five strains that were used to construct the graph and the other seven strains (Figure S13). Only the number of alignments with perfect identity is substantially lower for the strains that were not included in the creation of the graphs (Figure S13a).

### Running time comparison between different tools for HG00514 as genotyped on the HGSVC dataset

**Table S7:** Compute resources required for analysis of sample HG00514 on the HGSVC dataset.

| Tool | Wall Time (m) | Cores | Nodes | Max Memory (G) |
| --- | --- | --- | --- | --- |
| **vg** |  |  |  |  |
| vg construction | 49 | 8 | 1 i3.8xlarge | 0.4 |
| xg index | 13 | 8 | 1 i3.8xlarge | 48 |
| snarls index | 23 | 1 | 50 i3.8xlarge | 17 |
| gcsa2 index | 792 | 16 | 1 i3.8xlarge | 45 |
| mapping | 177 | 32 | 50 r3.8xlarge | 32 |
| genotyping (pack + call) | 56 | 10 | 1 i3.4xlarge | 63 |
| **BayesTyper** | 90 | 24 | 1 i3.8xlarge | 36 |
| **bwa mem** | 240 | 32 | 1 i3.8xlarge | 14 |
| **Delly Genotyper** | 69 | 1 | 1 i3.8xlarge | 69 |
| **SVTyper** | 477 | 1 | 1 i3.8xlarge | 0.7 |
| **Paragraph** | 76 | 32 | 1 i3.8xlarge | 5.9 |

SMRT-SV v2 Genotyper required roughly 36 hours and 30G ram on 30 cores to genotype the three HGSVC samples on the “SVPOP” VCF. These numbers are not directly comparable to the above table because 1) they apply to the “SVPOP” rather than “HGSVC” dataset (upon which we were unable to run SMRT-SV v2 Genotyper) and 2) we were unable to install SMRT-SV v2 Genotyper on AWS nodes and ran it on an older, shared server at UCSC instead.

Delly Genotyper, SVTyper and Paragraph start from a set of aligned reads, hence we also show the running time for read alignment with `bwa mem` [26].

For BayesTyper, the numbers include both khmer counting with kmc and genotyping. We note that BayesTyper integrated variant calls from GATK haplotypecaller[39] and Platypus[40], derived from reads mapped with bwa mem[26]. The numbers shown for BayesTyper does not include this variant discovery pipeline.

Note: toil-vg reserves 200G memory by default for vg snarls. For this graph, about an order of magnitude less was required. It could have been run on 10 cores on 5 nodes instead.

